# A superfolding sfPepper enables superresolution and single-molecule RNA imaging in live cells

**DOI:** 10.64898/2026.07.15.738587

**Authors:** Xin Jiang, Huiwen Li, Li Jiang, Chenxin Cao, Qi Jiang, Ni Su, Mengyue Fang, Zhengda Chen, Fangting Zuo, Yuzheng Zhao, Xiwei Tian, Xianjun Chen, Linyong Zhu, Yi Yang

## Abstract

Fluorescent RNAs (FRs) have emerged as powerful tools for the visualization of RNA in live cells. However, the poor folding of FRs is a major limiting factor in many advanced RNA imaging applications. Here, we describe a novel strategy to significantly increase the folding efficiency of FRs. By introducing a stabilizing stem and performing targeted mutagenesis at key junction regions, we develop a ‘superfolder’ variant of the Pepper aptamer, termed sfPepper, that exhibits a striking twofold increase in folding efficiency compared with Pepper under physiological conditions. sfPepper exhibits much improved thermostability and reduced ion dependence and, most importantly, is substantially brighter than Pepper in live cells. sfPepper enables robust imaging of diverse noncoding RNAs and messenger RNAs with increased signal-to-background ratios and stimulated emission depletion (STED) superresolution imaging of CUG trinucleotide repeat-containing ‘toxic RNAs’. Remarkably, only four sfPepper repeats facilitate sensitive single-molecule mRNA tracking, revealing dynamic heterogeneity among ER-associated mRNA molecules. Together, this study establishes an efficient strategy for improving FR folding and offers a powerful tool for fluorescently labelling RNAs in live cells.

## INTRODUCTION

RNA is among the most versatile biomolecules that not only functions to transfer genetic information between DNA and proteins but also is involved in the regulation of genome organization and gene expression ^1^. RNA localization is dynamically regulated in cells, but how RNA localization and dynamics coordinate with their biological function, particularly for numerous noncoding RNAs (ncRNAs), is largely unknown ^1, 2^. Therefore, developing genetically encoded fluorescent tags mimicking fluorescent proteins (FPs) to enable real-time imaging of RNA in live cells is a primary goal. Fluorescent RNAs (FRs), which consist of RNA aptamers that specifically bind to their cognate fluorogenic dyes, are promising tools for visualizing RNA dynamics inside cells ^3, 4^. Compared with traditional RNA tethering-based approaches, FRs possess the advantages of straightforward labelling, an increased signal-to-background ratio (SBR) and reduced interference with the target RNA ^2, 5^.

During the past two decades, scientists have been putting much effort and thought into developing or optimizing FRs with improved photophysical properties for extending the spatial and temporal scales of RNA imaging. RNA aptamers in FRs are typically generated using in vitro selection methods, such as SELEX (Systematic Evolution of Ligands by Exponential Enrichment). One major challenge is that RNA aptamers are not well folded either in vitro or in cells. The folding in cells is particularly difficult to achieve because selection methods typically rely on in vitro conditions that cannot accurately mimic the the complex intracellular environments ^6^. To improve aptamer folding, highly structured RNA scaffolds, e.g., tRNA, F30 and RNA origami, have been developed to facilitate the folding of the inserted aptamers ^4, 7, 8^. However, appending aptamers with scaffolds increases the overall size of the RNA tag, thereby increasing the possibility of interference with the target RNA. Moreover, scaffold-induced cleavage and instability of the RNA construct negates the ability of the scaffold to increase the amount of its correctly folded appended target RNA in live cells ^7^. These issues prevent the widespread use of RNA scaffolds for improving the cellular performance of FRs. Alternatively, aptamer folding can be enhanced by aptamer mutagenesis. In 2015, Strack et al pioneered a rational mutagenesis-based strategy to improve aptamer folding guided by fluorescence and thermostability. Specifically, they developed Spinach2, a ‘superfolder’ variant of Spinach that is markedly brighter than Spinach in live cells because of substantially improved folding ^9^. This pioneering work strongly supports the importance of aptamer folding on the intracellular performance of FRs. Nevertheless, Spinach2 remains the only reported example in which rational mutagenesis has substantially improved the folding of an FR, yet even only 58% of Spinach2 is properly folded under cytoplasmic-mimicking conditions. As a result, optimization of FRs with high folding efficiency remains a significant challenge.

We previously developed Pepper, a 43-nt RNA aptamer that binds the fluorogenic dye (4-((2-hydroxyethyl)(methyl)amino)-benzylidene)-cyanophenylacetonitrile (HBC) and its analogues to increase their fluorescence by up to 10,000-fold. Even compared with most currently available FRs, Pepper still shows significantly increased stability and cellular fluorescence, as well as broader spectral tunability from cyan to red ^10, 11^. These outstanding properties have established Pepper as a versatile tool for numerous in vitro and in vivo applications, including live-cell RNA imaging ^10, 12-21^, genomic loci labelling ^10, 22^ and biosensing ^23-35^. Although Pepper appears simple and promising, it folds inefficiently in physiological environments, which substantially reduces its fluorescence within cells and prevents its utility in advanced RNA imaging applications.

Here, we report the rational design and engineering of a ‘superfolder’ variant of Pepper, termed sfPepper, using a novel structure-guided strategy that combines stabilizing stem insertion and targeted mutagenesis at key junction regions. Whereas only 37% of Pepper folds correctly under cytoplasmic-mimicking conditions, sfPepper exhibits markedly improved folding efficiency, with 72% achieving proper folding under the same conditions. sfPepper enables advanced imaging of diverse noncoding RNAs and messenger RNAs, as well as superresolution RNA imaging and single-molecule RNA tracking. Together, this study not only offers a uniquely powerful tool for dissecting RNA localization and dynamics but also establishes a novel strategy for optimizing RNA aptamers for broad applications in imaging, biosensing, and therapeutics.

## RESULTS

### Development of sfPepper

FRs must fold correctly to adopt their proper tertiary structures, thereby enabling efficient binding to their cognate fluorogenic dyes and triggering the activation of fluorescence. To measure the fraction of Pepper that is properly folded, we used fluorescence correlation spectroscopy (FCS) to determine the molar fraction of fluorophore-occupied Pepper. Using buffers that mimic cytoplasmic ion concentrations, we found that only 37 ± 4.2% of Pepper is properly folded **(Fig. 1a and b)**. These findings led us to hypothesize that increasing the folding efficiency of Pepper in live cells would significantly improve its performance in cellular applications.

**Figure 1.**
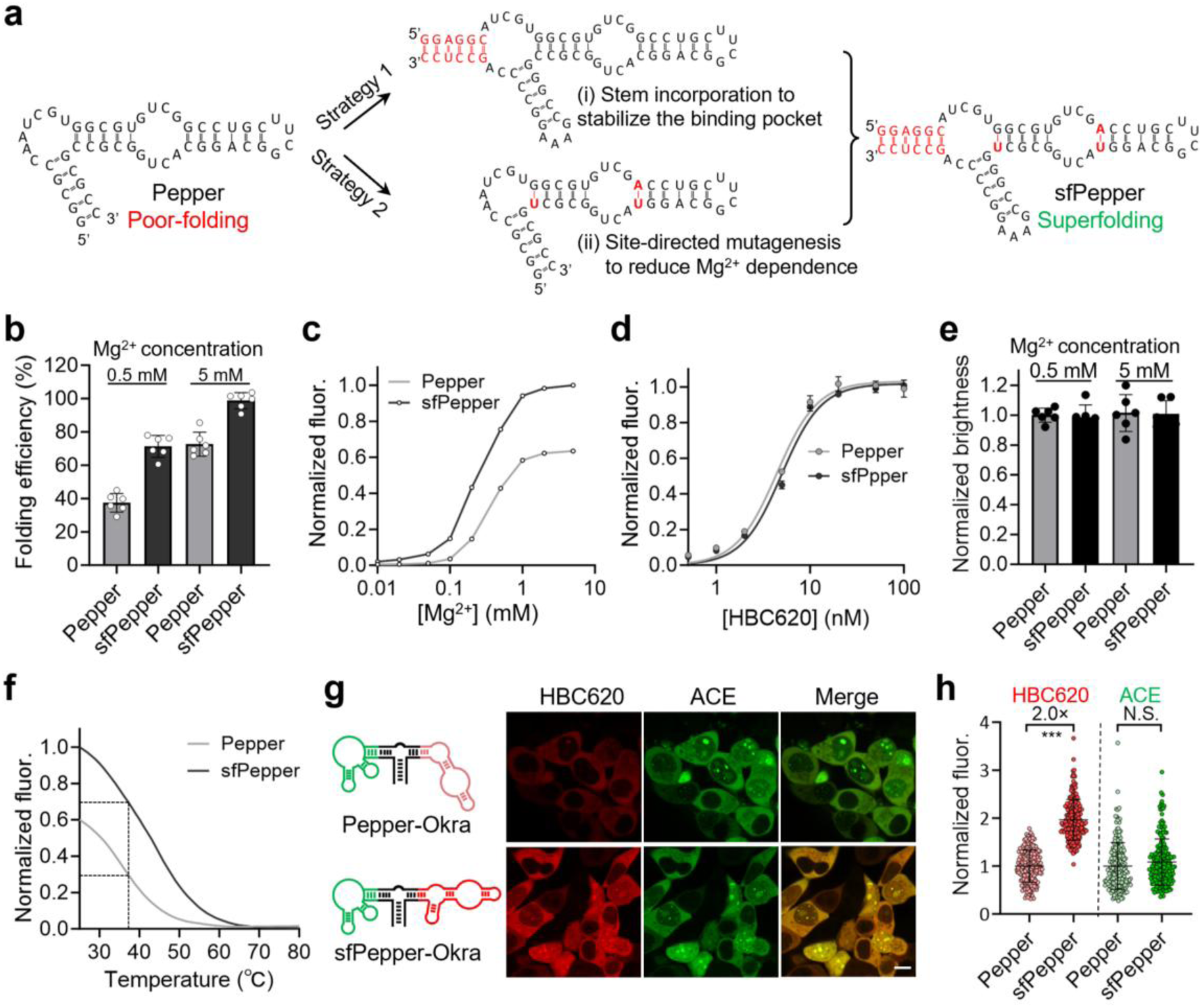
Development of sfPepper. **(a)** Flowchart of the RNA engineering strategy leading to the development of sfPepper. **(b)** Folding efficiencies of Pepper and sfPepper in the presence of 0.5 or 5 mM Mg²⁺. The data represent the means ± SDs from six biologically independent replicates. **(c)** Mg²⁺ dependence of Pepper and sfPepper in the presence of HBC620. **(d)** Binding affinity of HBC620 for Pepper or sfPepper. The fluorescence of the RNA-dye complexes in the presence of increasing concentrations of HBC620 were measured. The data represent the means ± SDs from three biologically independent replicates. **(e)** Fluorescence of a single molecule of Pepper or sfPepper in the presence of 0.5 or 5 mM Mg²⁺. The data represent the means ± SDs from six biologically independent replicates. **(f)** Thermostability of Pepper and sfPepper measured from 25°C to 80°C. Dashed lines indicate the residual fluorescence at physiological temperature (37 °C). **(g)** Confocal images of live HEK293T cells expressing circular Pepper-Okra or sfPepper-Okra incubated with 0.5 µM HBC620 and ACE. Scale bar, 10 μm. **(h)** The normalized HBC620 and ACE fluorescence intensities of the cells in **g**. Statistical comparisons were performed by two-tailed *t* tests. *\*\*\*P* < 0.001. N.S., no significant difference. The data represent the means ± SDs (*N* = 200 cells).

The three-dimensional architecture of RNA molecules is largely governed by the structural conformations of key secondary-structure elements, particularly pseudoknots, kissing loops, and bulge junctions ^36, 37^. Structural analysis of the Pepper-HBC complex revealed that the protruding J3/2 bulge junction forms a lid-like architecture over the HBC binding pocket. Although the J3/2 nucleotides do not interact with HBC directly, they play a crucial structural role in shaping the HBC binding pocket through stacking and hydrogen bonding interactions **(Supplementary Fig. 1a)** ^11^. Notably, deletion mutagenesis studies have also confirmed the essential role of J3/2, as its removal nearly completely abolishes the HBC binding capacity ^11^. These structural insights inspired us to investigate whether structural modification of J3/2 could increase its stability and consequently improve the cellular performance of Pepper. To this end, we first performed targeted nucleotide insertions or deletions in J3/2 to either extend or truncate the lid-like structure **(Supplementary Fig. 1a)**. Unfortunately, the majority of the obtained variants lost the ability to activate HBC620 fluorescence **(Supplementary Fig. 1b)**. We next engineered a stabilizing stem within the J3/2 region with the aim of reinforcing the lid-like structure. Through the systematic insertion of a six base pair stem at different positions in J3/2 **(Supplementary Fig. 2a)**, we found that three of the six variants, namely, M10, M11 and M13, exhibited increased fluorescence intensity compared with that of the wild-type Pepper **(Supplementary Fig. 2b)**. We further investigated the live-cell imaging performance of these three variants by expressing them in HEK293T cells from a U6 promoter using the Tornado system ^38^. Confocal microscopy revealed that compared with Pepper, M11 strikingly displayed 1.6-fold greater fluorescence intensity in cells **(Supplementary Fig. 3a and b)**.

Another key determinant of the overall tertiary architecture of RNA is magnesium ions (Mg²⁺) ^36, 37^. Previous studies have demonstrated that several hydrated Mg²⁺ ions bound at the J3/2 and J2/1 junction regions play critical roles in stabilizing the tertiary structure of the Pepper aptamer **(Supplementary Fig. 4a)**, thereby rendering the fluorescence of the Pepper– HBC complex highly Mg²⁺ dependent ^11^. However, this strong dependence on Mg²⁺ poses a significant limitation for RNA imaging in mammalian cells, where the estimated free intracellular Mg²⁺ concentrations range from only 0.2 to 1 mM ^39, 40^. To overcome this limitation, we introduced targeted mutations at these conformational junctions, specifically, the nucleotide pairs bridging J3/2 and P2, J2/1 and the P1 stem, with the aim of reducing the Mg²⁺ dependence of Pepper without compromising its structural integrity **(Supplementary Fig. 4b)**. Compared with Pepper, the M19 variant, in which a G-U pair bridged J3/2 and P2, exhibited markedly reduced Mg²⁺ dependence **(Supplementary Fig. 4c)**. In addition, the variants, M20, M21, and M23, containing a G–C, U–A, or U–G pair bridging junction J2/1 and the P1 stem, respectively, displayed modest decreases in Mg²⁺ dependence **(Supplementary Fig. 4c)**. We then combined these mutations and introduced them into the M11 variant **(Supplementary Fig. 5a)**. As predicted, the resulting variants, M24, M25 and M26, exhibited a further increase in intracellular fluorescence relative to that of M11 **(Supplementary Fig. 5b and c)**. Among them, M25 exhibited the most intense fluorescence and was therefore selected for subsequent studies.

Compared with Pepper, M25 displayed a markedly reduced dependence on Mg²⁺, with an EC₅₀ of 0.19 mM versus 0.35 mM for Pepper **(Fig. 1c)**. More importantly, under 0.5 mM Mg²⁺, sfPepper displayed nearly twice the fluorescence intensity of Pepper **(Fig. 1c)**. Further studies revealed that under Mg²⁺-saturated (5 mM) conditions, 98.7% of M25 folded properly, which is significantly greater than the 71.3% observed for Pepper **(Fig. 1b)**. Notably, at a physiologically relevant concentration of Mg²⁺ (0.5 mM), 72.6% of M25 remained correctly folded, a folding efficiency nearly twice that observed for Pepper under the same conditions **(Fig. 1b)**. Therefore, we refer to this ‘<u>s</u>uperfolder’ variant of <u>Pepper</u> as sfPepper. Upon binding to HBC analogues, sfPepper displayed excitation and emission spectra that were identical to those of Pepper **(Supplementary Fig. 6a)** while maintaining comparably high binding affinity **(Fig. 1d, Supplementary Fig. 6b and e)**. Notably, the single-molecule fluorescence intensity of sfPepper was comparable to that of Pepper **(Fig. 1e)**, and this similarity in fluorescence was also observed when excess RNA aptamer was used **(Supplementary Fig. 7a)**. These data suggest that the enhanced brightness of sfPepper reflects an increase in its folding efficiency, rather than an increase in the quantum yield or extinction coefficient of the RNA-fluorophore complex. Additionally, compared with Pepper, sfPepper also exhibited markedly improved thermostability in buffer containing a concentration of magnesium ions that mimics that of the cytoplasm, with a T_m_ of 43°C versus a T_m_ of 37°C for Pepper, which resulted in sfPepper exhibiting >2-fold greater fluorescence intensity than that of Pepper at 37 °C **(Fig. 1f)**. We also compared the folding kinetics between sfPepper and Pepper using a fluorescence recovery assay. The results revealed that compared with Pepper, sfPepper exhibited much faster fluorescence recovery kinetics following denaturation at 80°C and subsequent gradual cooling to 20°C, indicating that sfPepper folds more rapidly than Pepper **(Supplementary Fig. 7b)**. Additionally, sfPepper showed greater tolerance to denaturing reagents, including urea and guanidine hydrochloride (GdnHCl), as well as to pH fluctuations **(Supplementary Fig.7c-e)**, highlighting its superior folding efficiency under diverse environmental conditions.

### Cellular performance of sfPepper

We next tested the performance of sfPepper in live mammalian cells. Confocal images of HEK293T cells expressing circular sfPepper-Okra or Pepper-Okra were acquired in the presence of HBC620 and ACE dyes. In this assay, the fluorescence of the Okra:ACE, which was detected in a separate green channel, was used to monitor the expression levels of the RNA aptamer in individual cells. Compared with Pepper, sfPepper displayed a 2-fold increase in fluorescence intensity when the cells were incubated with HBC620 **(Fig. 1g and h)**. Given that the fluorescence of Okra:ACE was comparable **(Fig. 1h)**, the much more intense fluorescence of sfPepper was attributed to its improved folding but not increased expression. Similar results were observed in cells expressing the linearized RNA aptamer alone or embedded in a transfer RNA (tRNA) scaffold **(Supplementary Fig. 8)**. Moreover, sfPepper was much brighter than Pepper when the cells were incubated with HBC analogues, including HBC530 and HBC485 **(Supplementary Fig. 6c–d and f-g)**. We then tested whether sfPepper-tagged functional RNAs exhibited more intense fluorescence than Pepper-tagged RNAs in mammalian cells. HEK293T cells expressing sfPepper or Pepper-tagged 5S RNA exhibited the expected diffuse cytoplasmic fluorescence, but compared with 5S-Pepper, 5S-sfPepper presented 2.3-fold stronger fluorescence in these cells (**Supplementary Fig. 9a and b**). Additionally, sfPepper-tagged U6 small nuclear RNA colocalized with GFP-tagged SART3, a known protein component of Cajal bodies, in the nucleus ^41^, and its fluorescence was also much intense than that of Pepper-tagged U6 (**Supplementary Fig. 9c and d**).

We further attempted to use sfPepper to image messenger RNA (mRNA) transcribed by RNA polymerase II. Considering that the abundance of mRNAs is much lower than that of the numerous ncRNAs transcribed by RNA polymerase III ^10^, we used the tandem repeats of sfPepper to amplify the fluorescence signals. We created a tandem array containing four repeats of sfPepper, termed 4sfPepper, by fusing one molecule of sfPepper to the terminal stem loop of another molecule of sfPepper **(Supplementary Fig. 10a)**. When 4sfPepper was fused to the 3’ untranslated region (UTR) of BFP and expressed in COS-7 cells, we observed the typical cytosolic red fluorescence of *BFP* mRNA and the blue fluorescence of the BFP protein **(Supplementary Fig. 10b)**. Compared with cells expressing 4Pepper-tagged *BFP* mRNA, cells expressing 4sfPepper-tagged *BFP* mRNA presented substantially greater red fluorescence intensity **(Supplementary Fig. 10b and c)**. Notably, 4sfPepper- and 4Pepper-tagged *BFP* mRNA yielded expression levels of the BFP protein comparable to the of untagged *BFP* mRNA **(Supplementary Fig. 10d)**, indicating that labelling with 4sfPepper had no effect on the translation of its tagged mRNA. Further studies revealed that the recruitment of *ACTB* mRNA to stress granules (SGs) under stress conditions could also be visualized by labelling *ACTB* mRNA with 4sfPepper, which was reflected by the accumulation of mRNA fluorescence in SGs enriched in Ras GTPase-activating protein binding protein 1 (G3BP1), a well-known protein marker of SGs ^42^ **(Supplementary Fig. 11)**. Collectively, these data demonstrate that sfPepper enables enhanced RNA visualization in live cells while maintaining negligible perturbations to the inherent localization patterns of the target RNAs.

### Comparison of sfPepper to other FRs

We then compared sfPepper:HBC620 with existing spectrally matched monomeric FRs, including Clivia:NBSI624 ^22^, RhoBAST:TMR-DN ^43^, RhoBAST:SpyRho ^44^, and Red Broccoli:OBI ^45^. HEK293T cells expressing RNA aptamers without a scaffold RNA from the U6 promoter were imaged in the presence of their corresponding fluorogenic dyes. The results showed that sfPepper:HBC620 outperformed the other red FRs in cells in terms of both fluorescence intensity and signal-to-background ratio **(Fig. 2a and b)**. Specifically, compared with Clivia:NBSI624, RhoBAST:SpyRho, RhoBAST:TMR-DN and Red Broccoli:OBI, sfPepper:HBC620 exhibited 5.4-, 1.6-, 3.7- and 21.4-fold greater intracellular fluorescence intensity, respectively **(Fig. 2a and b)**. Combined with the low background fluorescence of the HBC620 dye, sfPepper:HBC620 achieved a remarkably high signal-to-background ratio of 15.9-fold, whereas the remaining FRs presented signal-to-background ratios of < 4-fold **(Fig. 2a and b)**. In addition, sfPepper:HBC620 significantly outperformed the other FRs when the RNA aptamers were embedded in a tRNA scaffold **(Supplementary Fig. 12)**. Notably, the tRNA-scaffolded RhoBAST aptamer displayed markedly enhanced cellular performance compared to its unscaffolded counterpart upon incubation with either TMR-DN or SpyRho dye **(Fig. 2a and b, Supplementary Fig. 12)**. These results suggest that the RhoBAST alone folds inefficiently in the cellular environment and likely requires an RNA scaffold to promote its proper folding.

**Figure 2.**
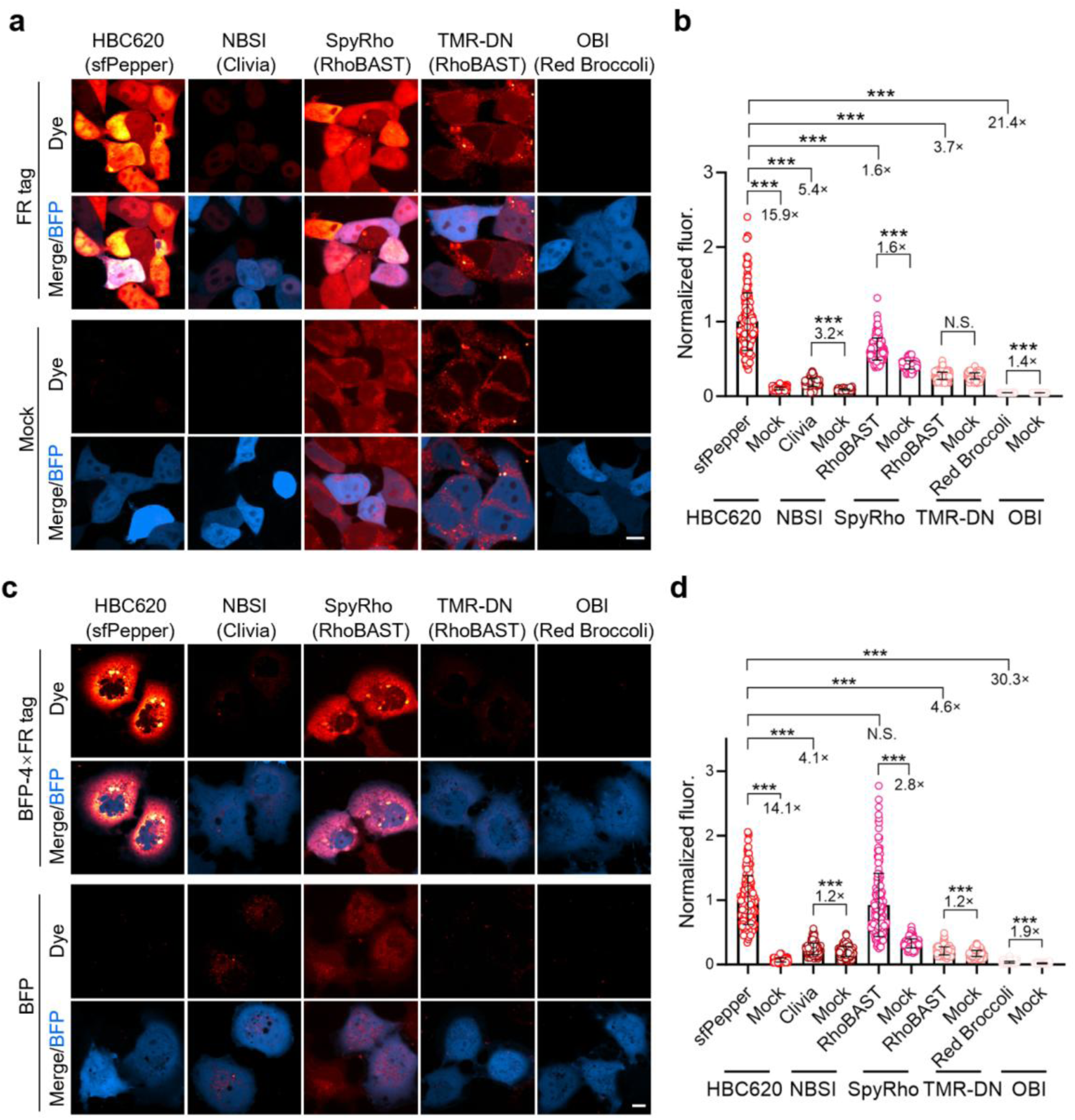
Comparison of sfPepper to other high-performance FRs. **(a)** Confocal images of live HEK293T cells expressing sfPepper, Clivia, RhoBAST or Red Broccoli without scaffold RNA. Fluorescence was visualized upon incubation with 0.5 µM HBC620, NBSI, SpyRho, TMR-DN, or OBI. Control cells were transfected with an untagged BFP plasmid and incubated with the respective dyes. Scale bar, 10 µm. **(b)** Fluorescence intensities of the cells in **a**. **(c)** Confocal images of live COS-7 cells expressing BFP mRNA that was labelled with four copies of sfPepper, Clivia, RhoBAST or Red Broccoli. Fluorescence was visualized upon incubation with 0.2 µM HBC620, NBSI, SpyRho, TMR-DN, or OBI, respectively. Control cells were transfected with an untagged BFP expression plasmid and incubated with the respective dyes. Scale bar, 10 µm. **(d)** Fluorescence intensities of the cells in **c**. Statistical comparisons in **b** and **d** were performed by two-tailed *t* tests. *\*\*\*P* < 0.001. N.S., no significant difference. The data represent the means ± SDs (*N* = 200 cells).

To compare the performance of sfPepper with that of other FRs for live-cell mRNA visualization, we fused four repeats of different RNA aptamers to the 3’ UTR of *BFP* mRNA, and cells expressing these chimeric mRNAs from the CMV promoter were imaged in the presence of their corresponding fluorogenic dyes. Compared with control cells expressing untagged *BFP* mRNA, only cells expressing 4sfPepper- or 4RhoBAST-tagged *BFP* mRNA exhibited significantly more intense red fluorescence signals **(Fig. 2c and d)**. Owing to the substantial reduction in background fluorescence of 4sfPepper, 4sfPepper achieved a signal-to-background ratio of 14.1-fold, which was much greater than 2.8-fold ratio observed for 4RhoBAST **(Fig. 2c and d)**. In comparison, the NBSI624, TMR-DN, and OBI dyes were hardly capable of visualizing 4Clivia-, 4RhoBAST- and 4Broccoli-tagged mRNA, respectively **(Fig. 2c and d)**. Thus, compared with previously reported FRs, sfPepper showed outstanding robustness for labelling mRNA.

### Imaging of CUG-repeat RNA beyond the diffraction limit

Many ‘toxic RNAs’ contain extensive trinucleotide repeats ^46^. Myotonic dystrophy type 1 (DM1) is a dominantly inherited disorder that results from a toxic gain of function of RNA transcripts containing expanded CUG repeats in the 3’UTR of DMPK. In patients with DM1, RNA with CUG repeats folds into a hairpin structure with an array of repeating 1 × 1-nucleotide UU internal loops to form intranuclear aggregates ^47^. In this study, we evaluated whether sfPepper could be used for in situ labelling and imaging of CUG repeat-containing RNA in live cells. To this end, we fused 4sfPepper to 960-repeat CUG trinucleotide-containing mRNA for expression in HEK293T cells under the CMV promoter **(Fig. 3a and b)** ^48, 49^. Upon incubation with HBC620 dye, we readily detected bright nuclear foci in cells expressing *Exon_11-15_-CUG_960_-4sfPepper* mRNA **(Fig. 3c)**, which colocalized perfectly with the fluorescence signal from SNAP-MBNL1, a marker of CUG repeat-containing nuclear foci ^50^ **(Fig. 3d)**. Notably, compared with those labelled with four repeats of Pepper or RhoBAST, sfPepper-labelled nuclear foci exhibited substantially more intense fluorescence **(Supplementary Fig. 13)**, highlighting the advantages of sfPepper for imaging CUG repeat-containing mRNA aggregates in live mammalian cells.

**Figure 3.**
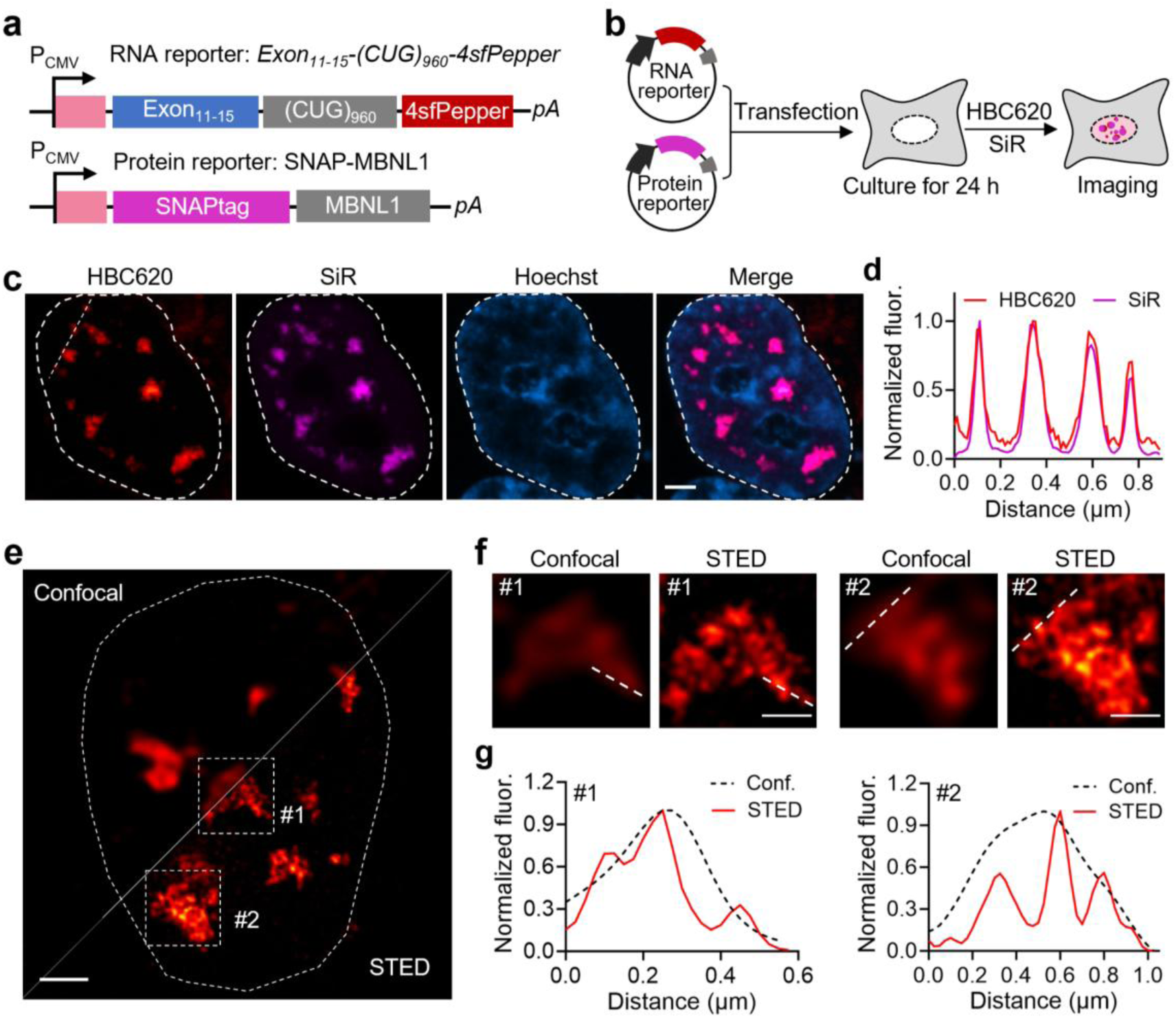
STED imaging of CUG-repeat RNA in live cells. **(a)** Schematic illustration of the constructs used for expressing sfPepper-tagged CUG repeat-containing mRNA. **(b)** Illustration of the procedure for imaging sfPepper-tagged CUG repeat-containing mRNAs. **(c)** Confocal images of HEK293T cells expressing *Exon_11-15_-CUG_960_-4sfPepper* mRNA and the SNAP-MBNL1 fusion protein incubated with HBC620 and SiR for 30 min prior to imaging. Scale bar, 3 µm. **(d)** Normalized intensity profiles along the white line in **c**. **(e)** Combined confocal and STED image of a nucleus in a live HEK293T cell expressing *Exon_11-15_-CUG_960_-4sfPepper* mRNA after incubation with HBC620 for 30 min prior to imaging. Scale bar, 2 µm. **(f)** Magnified views of the regions marked by the white squares in **(e)**. Scale bars, 500 nm. **(g)** Normalized intensity profiles along the white line in **f**.

Inspired by above results, we next tested the usefulness of sfPepper for RNA imaging beyond the diffraction limit in stimulated emission depletion (STED) microscopy. To this end, we performed superresolution imaging of *Exon_11-15_-CUG_960_-4sfPepper* mRNA aggregates in live HEK293T cells using STED microscopy **(Fig. 3e–g)**. Compared with confocal microscopy, live-cell STED microscopy allowed us to resolve more architectural details of RNA aggregates containing CUG repeats that cause disease **(Fig. 3e–g)**, which, to our knowledge, has not been reported previously. Intriguingly, these aggregates were highly heterogeneous in appearance and displayed irregular alveolate-like structures **(Fig. 3e–g)**, details that were hardly visible in the confocal images.

### Single-molecule RNA imaging using sfPepper

Single-molecule fluorescence imaging technology has enabled the assessment of the behaviours of individual RNA molecules in live cells in real time, revealing not only the temporal rules but also the spatial coordination of underlying molecular interactions during various biological activities ^51^. To determine whether sfPepper could be used for single-molecule RNA imaging, we created a construct to transcribe *Srprb-BFP* mRNA fused with 4 repeats of 4sfPepper and 16 repeats of the MS2 array in the 3’ UTR **(Fig. 4a and b)**, which allowed us to simultaneously visualize single mRNA molecules with the sfPepper and MS2–MCP system, a gold standard for single-molecule RNA tracking in live cells ^52-54^. In *Srprb-BFP-4×4sfPepper-16×MS2* mRNA, the signal peptide of the encoded signal recognition particle receptor B subunit (Srprb) contains the endoplasmic reticulum (ER)-targeted localization sequence of the fusion protein and can also tether the mRNA molecules to the outer ER membrane during protein translation through cotranslational membrane insertion ^52^ **(Supplementary Fig. 14a)**. COS-7 cells coexpressing *Srprb-BFP-4×4sfPepper-16×MS2* mRNA and tdMCP-mStayGold protein were incubated with HBC620 cells and imaged. As expected, the images revealed markedly excellent colocalization of the mStayGold and HBC620 signals on the ER **(Supplementary Fig. 14b)**, suggesting that labelling with sfPepper does not affect the localization of individual mRNA molecules.

**Figure 4.**
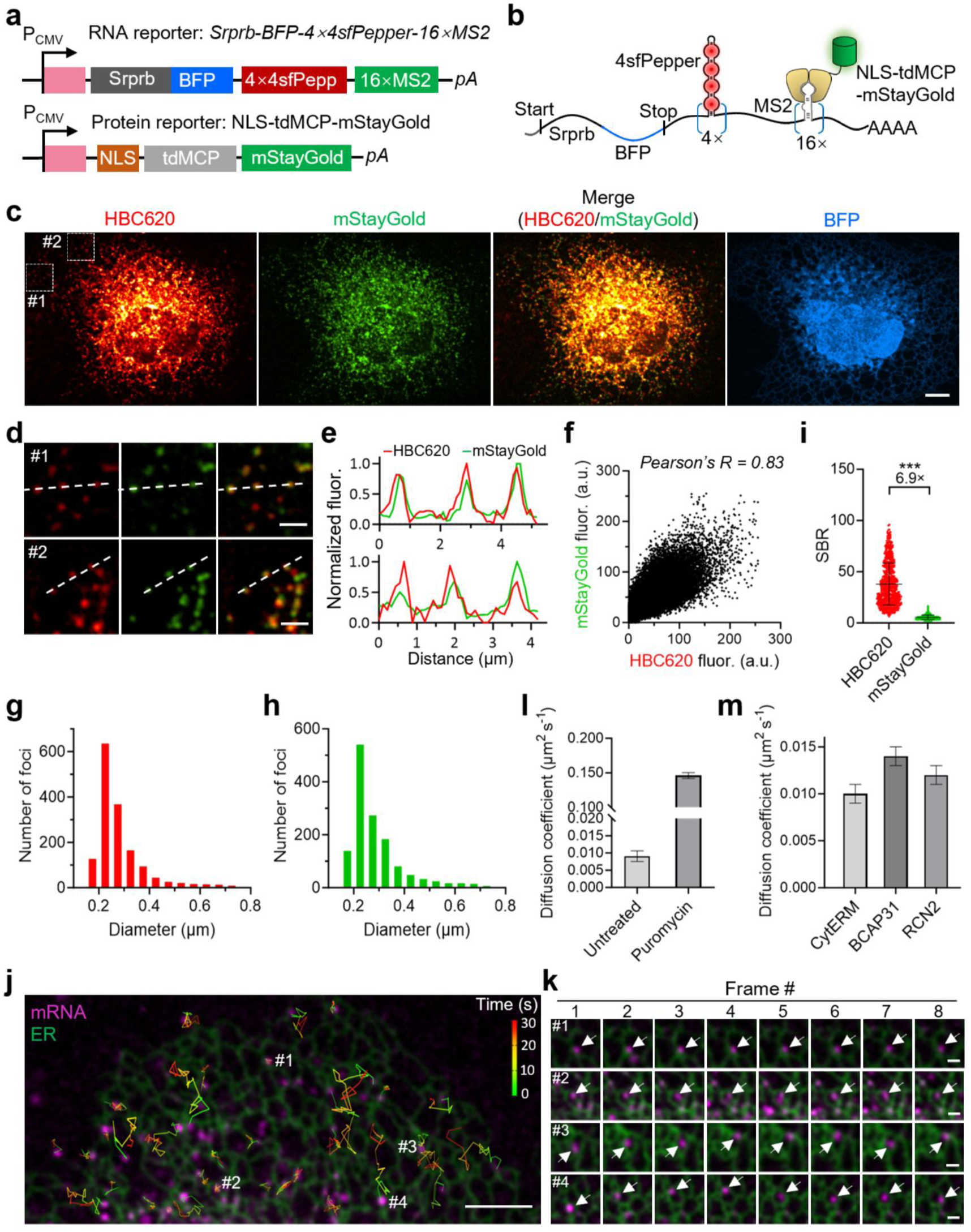
Single-molecule RNA imaging with sfPepper in live cells. **(a)** Constructs used to express 4×4sfPepper- and 16×MS2-tagged *Srprb-BFP* mRNA, which can be tethered by the fusion protein of the tandem dimer of MCP (tdMCP) and mStayGold. **(b)** Illustration of *Srprb-BFP* mRNA tagged with 4×4sfPepper and 16×MS2. **(c)** Confocal images of live COS-7 cells expressing low amounts of *Srprb-BFP-4×4sfPepper-16×MS2* mRNA and tdMCP-mStayGold upon incubation with 100 nM HBC620. Scale bar, 10 µm. **(d)** Magnified views of the regions indicated in the white boxes in **c** showing the colocalization of red and green foci. Scale bars, 3 µm. **(e)** Line profiles of the fluorescence from the HBC620 and mStayGold channels of the dashed lines in **(d)**. **(f)** Colocalization analysis of the cytosolic HBC620 and mStayGold signals in **c** (*N* = 10,797 foci, Pearson’s *R* = 0.83). **(g, h)** Analysis of the diameters of the foci (*N* > 1,300 foci) in the HBC620 **(g)** and mStayGold **(h)** channels in **(c)**. **(i)** Mean signal-to-background ratios of the foci in **(c)** from the HBC620 and mStayGold channels (HBC620: *N* = 672 foci; mStayGold: *N* = 666 foci). Statistical comparison was performed by a two-tailed *t* test. ***P < 0.001. The data represent the means ± SDs (HBC620: *N* = 672 foci; mStayGold: *N* = 666 foci). **(j)** Representative live-cell image of a COS-7 cell expressing *Srprb-mStayGold-4×4sfPepper* mRNA (enlarged view of the white box indicated region from **Supplementary** Fig. 18). Overlay of the mRNA channel (sfPepper:HBC620, purple) with the ER channel (Srprb-mStayGold, green). The trajectories of single RNA tracks at different time points are shown in the figure. Scale bar, 5 µm. **(k)** Magnified views of the indicated regions in **j** showing single RNAs during the first eight frames. # 1 and 2 represent mRNA molecules that appeared static on the ER surface; # 3 and 3 represent mRNA molecules that dynamically and stochastically associated with and dissociated from the ER membrane. Scale bars, 1 µm. **(l)** Diffusion coefficients of Srprb mRNA molecules without or with puromycin treatment. Data represent the means ± SDs (-puromycin: *N* = 46 foci; + puromycin: *N* = 46 foci). **(m)** Diffusion coefficients of different ER-localized mRNA species. The data represent the means ± SDs (*CytERM*: *N* = 35 foci; *BCAP31*: *N* = 50 foci; *RCN2*: *N* = 33 foci).

When the expressions of *Srprb-BFP-4×4sfPepper-16×MS2* mRNA were low, we observed individual diffraction-limited red fluorescent puncta that colocalized well with the green fluorescent puncta from the MS2-MCP system (Pearson’s *R* of 0.83) **(Fig. 4c–f)**. Analysis of the fluorescence intensities and the size distributions of the fluorescent puncta in the sfPepper and mStayGold channels revealed a single population of particles with a mean diffraction-limited diameter of ∼250–300 nm **(Fig. 4g and h)**, confirming that the fluorescent puncta in the images indeed represent single molecules. Additionally, sfPepper exhibited a remarkably high signal-to-background ratio of 37.9-fold, which was 6.9 times greater than that of the MS2-MCP system (5.5-fold) **(Fig. 4i)**. Surprisingly, single-molecule RNA imaging was achieved even when a single copy of the 4sfPepper tandem repeat was used **(Supplementary Fig. 15)**, in stark contrast to previous studies that required 16 or 24 copies of FR repeats for comparable resolution ^55, 56^. Although 4sfPepper-tagged mRNA puncta displayed a 3.1-fold lower fluorescence intensity than that of 4×4sfPepper-tagged mRNA puncta, the signal-to-background ratio was still superior to that of the MS2-MCP system, despite containing fewer fluorophores (4 for HBC620 versus 8 for mStayGold) **(Supplementary Fig. 15i and 16)**. In contrast, in COS-7 cells expressing 4RhoBAST- or 4Pepper-tagged *Srprb* mRNA, we observed only blurry fluorescent puncta in the HBC620 and SpyRho channels, whereas the green fluorescent puncta in the mStayGold channel remained sharply resolved **(Supplementary Fig. 17)**.

To investigate the mobility of individual ER-localized mRNA transcripts, we performed single-particle tracking over 30 consecutive frames. Our results revealed that the majority of *Srprb-mStayGold-4×4sfPepper* mRNA molecules exhibited limited but heterogeneous tracks **(Supplementary Fig. 18)**. Specifically, a substantial fraction of the mRNA molecules dynamically and stochastically associated with and dissociated from the ER membrane.

Moreover, some of the mRNA molecules appeared static on the ER surface throughout the entire imaging period **(Fig. 4j and k)**, suggesting that these molecules were likely undergoing translation. By plotting the mean square displacement (MSD) versus the delay time, we calculated a diffusion coefficient (*D*) of 0.009 μm^2^s^-1^ *Srprb-mStayGold-4×4sfPepper* mRNA molecules **(Fig. 4l, Supplementary Fig. 19a)**, which agreed with previously reported diffusion coefficients for ER-localized mRNAs ^57^. Importantly, treatment with puromycin, which disrupts the ribosome and dissociates mRNA from the corresponding nascent peptide ^58^, generally increased the degree of mobility of the mRNA particles, which was also apparent upon visual inspection, consistent with the dissociation of the reporter mRNA from the ER **(Fig. 4l, Supplementary Fig. 19b)**. Using 4×4sfPepper, we found that different ER-localized mRNA species shared similar dynamic behaviours, despite slight variations in their mobility **(Fig. 4m, Supplementary Fig. 19c-e)**. Taken together, these data show that sfPepper enables the robust detection and tracking of single mRNA transcripts in live cells, thereby deepening our understanding of the diversity and dynamics of ER-localized mRNAs.

## DISCUSSION

RNA plays crucial roles in regulating diverse cellular functions. Techniques that enable real-time visualization of RNA localization and dynamics are essential for elucidating the diverse roles that RNAs play in cellular processes. The use of FRs, which are FP-like entities consisting of RNA aptamers that can bind and turn on otherwise nonfluorescent small-molecule dyes, provides an attractive approach for visualizing RNAs in live cells. Although dozens of FRs have been developed ^59, 60^, only a small number of them function robustly in mammalian cells. A one major reason is that most FRs are much less fluorescent than expected based on their in vitro fluorescence. In contrast, Pepper stands out owing to its superior cellular fluorescence and broad spectral tunability compared with most currently available FRs, enabling robust imaging of diverse RNA species. However, its inefficient folding under physiological conditions limits its performance in advanced live-cell RNA imaging and leaves substantial room for improvement.

As antibody mimics, aptamers were initially developed to specifically bind and inhibit protein function by folding into complex three-dimensional structures. Thus, their correct folding is pivotal for their functions. Although screening, mutagenesis and scaffolding approaches have been reported to improve aptamer folding, poor folding is still a major roadblock that prevents the widespread use of aptamers for imaging and therapeutic applications ^3^. Here, we report a novel structure-guided engineering strategy, which, to our knowledge is the first strategy that facilitates RNA folding by introducing a stem into the bulge junction of the Pepper aptamer. The resulting sfPepper not only exhibits a much greater fraction of properly folded aptamers but also displays faster folding kinetics. We believe that the advantage of faster folding will be more apparent when the aptamer is cotranscribed with flanking sequences, as a fast-folding rate can minimize misfolding between the aptamer and nascent flanking RNA sequences. Moreover, the substantial improvements in folding efficiency, thermostability, and cellular brightness of sfPepper make it particularly advantageous for advanced RNA imaging applications, including superresolution imaging of CUG repeat-containing RNAs and single-molecule RNA tracking. For the first time, we demonstrate that the spatial distribution of CUG repeat-containing RNAs is highly heterogeneous within nuclear aggregates. Moreover, we revealed that mRNA molecules associated with the ER surface display remarkably dynamic heterogeneity. Collectively, this work establishes a novel strategy to enhance FR folding and provides a valuable tool for real-time tracking and monitoring of RNA localization and dynamics in living cells.

Single-molecule RNA imaging has been realized by RNA tethering approaches, e.g., the well-known MS2-MCP system, which is the gold standard for single-molecule RNA tracking in live cells ^52-54^. In this system, multiple tandem repeats (24× or 48×) of the MS2 hairpin are fused to the 3’ UTR of its mRNA, which can recruit a fusion protein comprising a fluorescent protein fused to the MCP protein. To reduce cytoplasmic background fluorescence and improve the SBR, a nuclear localization sequence (NLS) is used, causing the excess unbound MCP–FP fusion protein to accumulate in the nucleus. Alternatively, RBP–FP fusions can be engineered to be fluorogenic, wherein they are unstable until they bind RNA motifs (e.g., RNA aptamers or stem loops) inserted into target RNAs, thus reducing the background fluorescence from unbound RBP–FP fusions ^53, 61-63^. In addition, Xia et al developed an approach that combines the RNA-targeting CRISPR–Csm complex with multiplexed guide RNAs for the visualization of unmodified endogenous RNAs at the single-molecule level ^64^. However, these protein-based approaches are suboptimal for functional studies of single RNAs owing to the bulky tag and potential effects on RNA behaviour caused by the tags themselves and heavy protein tethering ^56, 65-69^. Therefore, a robust protein-free approach is still highly desirable. FRs that consist of genetically encoded RNA aptamers and small-molecule fluorogenic dyes are attractive candidates for developing protein-free approaches for single-molecule RNA imaging. However, only a few FRs have been shown to be capable of single-molecule RNA imaging in live cells ^55, 56, 70^. Given the relatively low cellular fluorescence or imaging contrast of these FRs, at least 16 FR tag repeats are needed to amplify the fluorescence signals to achieve an adequate SBR, which increases the possibility of interference with the target RNA. In contrast, sfPepper has the advantages of superior cellular fluorescence and imaging contrast, allowing the imaging of single RNA molecules using a small RNA tag containing only four sfPepper repeats. In addition, unlike previous FRs that are incorporated into RNA scaffolds to promote aptamer folding via three- or four-way junctions, 4sfPepper comprises four sfPepper aptamers placed end-to-end without an additional scaffold, which further reduces the overall size of the tag and thus minimizes potential interference with RNA behaviour.

In summary, we developed a novel strategy to engineer a ‘superfolder’ variant of Pepper, which has excellent photophysical properties and enables advanced superresolution and single-molecule RNA imaging in live cells. This structural modification-based strategy may provide new ideas for the optimization of RNA aptamers with improved folding for imaging and therapeutic applications. In the future, we will be able to rapidly obtain a series of strongly fluorescent and stable FRs by aptamer and fluorogenic dye coevolution for use as advanced RNA imaging agents to explore the functions and mechanisms of RNAs in diverse biological processes.

## ACKNOWLEDGMENTS

We thank Ji Tang from Optofem company for technical assistance of STED imaging. This research was supported by the National Key Research and Development Program of China (2022YFC3400100, 2024YFA0917700), STI2030-Major Projects (2021ZD0202200 and 2021ZD0202203), NSFC (32121005, 22437001, 92581203 and 92357308, 32250009 and 22507071), the Shanghai Science and Technology Commission (23J21900400), the Shanghai Municipal Education Commission (2021 Sci & Tech 03-28), the Fundamental and Interdisciplinary Disciplines Breakthrough Plan of the Ministry of Education of China (JYB2025XDXM404), the Fundamental Research Funds for the Central Universities.

## AUTHOR CONTRIBUTIONS

Concepts were conceived by Y.Y., L.Z. and X.C.; Y.Y., L.Z., X.C., X.J., H.L., L.J. and C.C. designed the experiments and analyzed the data; L.J. synthesized the dyes; X.J., H.L., C.C. and Q.J. characterized the sfPepper *in vitro* and constructed plasmids; X.J. performed live cell imaging experiments. N.S., M.F., Z.C., F.Z., Y.Z., and X.T. gave technical support and conceptual advice. Y.Y., L.Z., X.C. and X.J. wrote the manuscript.

## REFERENCES

1. Morris, K.V. & Mattick, J.S. The rise of regulatory RNA. Nature reviews. Genetics 15, 423–437 (2014).

2. Braselmann, E., Rathbun, C., Richards, E.M. & Palmer, A.E. Illuminating RNA Biology: Tools for Imaging RNA in Live Mammalian Cells. Cell chemical biology 27, 891–903 (2020).

3. Babendure, J.R., Adams, S.R. & Tsien, R.Y. Aptamers switch on fluorescence of triphenylmethane dyes. Journal of the American Chemical Society 125, 14716–14717 (2003).

4. Paige, J.S., Wu, K.Y. & Jaffrey, S.R. RNA mimics of green fluorescent protein. Science 333, 642–646 (2011).

5. You, M. & Jaffrey, S.R. Structure and Mechanism of RNA Mimics of Green Fluorescent Protein. Annual review of biophysics 44, 187–206 (2015).

6. Hou, Q. & Jaffrey, S.R. Synthetic biology tools to promote the folding and function of RNA aptamers in mammalian cells. RNA biology 20, 198–206 (2023).

7. Filonov, G.S., Kam, C.W., Song, W. & Jaffrey, S.R. In-gel imaging of RNA processing using broccoli reveals optimal aptamer expression strategies. Chemistry & biology 22, 649–660 (2015).

8. Geary, C., Grossi, G., McRae, E.K.S., Rothemund, P.W.K. & Andersen, E.S. RNA origami design tools enable cotranscriptional folding of kilobase-sized nanoscaffolds. Nature chemistry 13, 549–558 (2021).

9. Strack, R.L., Disney, M.D. & Jaffrey, S.R. A superfolding Spinach2 reveals the dynamic nature of trinucleotide repeat-containing RNA. Nature methods 10, 1219–1224 (2013).

10. Chen, X. et al. Visualizing RNA dynamics in live cells with bright and stable fluorescent RNAs. Nature biotechnology 37, 1287–1293 (2019).

11. Huang, K. et al. Structure-based investigation of fluorogenic Pepper aptamer. Nature chemical biology 17, 1289–1295 (2021).

12. Bereiter, R. et al. Engineering covalent small molecule-RNA complexes in living cells. Nature chemical biology 21, 843–854 (2025).

13. Zheng, Y. et al. RNA polymerase stalling-derived genome instability underlies ribosomal antibiotic efficacy and resistance evolution. Nature communications 15, 6579 (2024).

14. Zheng, R. et al. Multiplexed sequential imaging in living cells with orthogonal fluorogenic RNA aptamer/dye pairs. Nucleic acids research 52, e67 (2024).

15. Wan, L. et al. Small CAG Repeat RNA Forms a Duplex Structure with Sticky Ends That Promote RNA Condensation. Journal of the American Chemical Society 147, 3813–3822 (2025).

16. Ji, R.Y. et al. RNA Condensate as a Versatile Platform for Improving Fluorogenic RNA Aptamer Properties and Cell Imaging. Journal of the American Chemical Society 146, 4402–4411 (2024).

17. Palacio, M. & Taatjes, D.J. Real-time visualization of reconstituted transcription reveals RNAPII activation mechanisms at single promoters. Cell reports 44, 116251 (2025).

18. Luyties, O. et al. Multi-omics and biochemical reconstitution reveal CDK7-dependent mechanisms controlling RNA polymerase II function at gene 5’- and 3’ ends. Cell reports 44, 115904 (2025).

19. Bai, R. et al. In Situ Assembly of Fluorogenic RNA for Screening Natural Anti-Liver Fibrosis Products via Dynamic Visualization of COL1A1 mRNA. Advanced science 12, e02850 (2025).

20. Pei, L. et al. Aggresomes protect mRNA under stress in Escherichia coli. Nature microbiology 10, 2323–2337 (2025).

21. Zuo, F. et al. Imaging the dynamics of messenger RNA with a bright and stable green fluorescent RNA. Nature chemical biology 20, 1272–1281 (2024).

22. Jiang, L. et al. Large Stokes shift fluorescent RNAs for dual-emission fluorescence and bioluminescence imaging in live cells. Nature methods 20, 1563–1572 (2023).

23. Fang, M. et al. Imaging intracellular metabolite and protein changes in live mammalian cells with bright fluorescent RNA-based genetically encoded sensors. Biosensors & bioelectronics 235, 115411 (2023).

24. Chen, Z. et al. Genetically encoded RNA-based sensors with Pepper fluorogenic aptamer. Nucleic acids research 51, 8322–8336 (2023).

25. Yuan, D. et al. Allosteric genetically encoded biosensor for spatiotemporal monitoring of endogenous RNA dynamics in living cells. Proceedings of the National Academy of Sciences of the United States of America 122, e2409309122 (2025).

26. Hou, J., Guo, P., Wang, J., Han, D. & Tan, W. Artificial dynamic structure ensemble-guided rational design of a universal RNA aptamer-based sensing tag. Proceedings of the National Academy of Sciences of the United States of America 121, e2414793121 (2024).

27. Wang, Q. et al. Inert Pepper aptamer-mediated endogenous mRNA recognition and imaging in living cells. Nucleic acids research 50, e84 (2022).

28. Yin, P. et al. A universal orthogonal imaging platform for living-cell RNA detection using fluorogenic RNA aptamers. Chemcial Science 14, 14131–14139 (2023).

29. Bodin, M.R. & Hammond, M.C. Visualizing intracellular glycine with two-dye and single-dye ratiometric RNA-based sensors. Nucleic acids research 53, gkaf839 (2025).

30. Spradlin, S.F., Dickerson, K.A. & Batey, R.T. Structural and functional clues challenge the hypothesis that the yjdF riboswitch is natively regulated through broad recognition of azaaromatic compounds. Nucleic acids research 53, gkaf1278 (2025).

31. Yang, J. et al. Programmable Fluorescent RNA with a Catalytic Hairpin Assembly System Enables Label-Free Monitoring of MicroRNA in Living Cells. Analytical chemistry 97, 24003–24010 (2025).

32. Ming, W. et al. Allele-Specific CRISPR-Cas9-Based Ratiometric Fluorescence Platform for Portable EGFR L858R Mutation Detection. Analytical chemistry 97, 25832–25839 (2025).

33. Gu, X. et al. Enhanced detection of HBV and HCV using Cas13a-FLAP and FGoAI platforms. Chemcial Science 17, 1656–1665 (2025).

34. Yan, Z. et al. Programmable fluorescent aptamer-based RNA switches for rapid identification of point mutations. Nature chemistry 17, 1826–1838 (2025).

35. Zhang, K. et al. Strategic base modifications refine RNA function and reduce CRISPR-Cas9 off-targets. Nucleic acids research 53, gkaf082 (2025).

36. Hermann, T. & Patel, D.J. RNA bulges as architectural and recognition motifs. Structure 8, R47–54 (2000).

37. Butcher, S.E. & Pyle, A.M. The molecular interactions that stabilize RNA tertiary structure: RNA motifs, patterns, and networks. Accounts of chemical research 44, 1302–1311 (2011).

38. Litke, J.L. & Jaffrey, S.R. Highly efficient expression of circular RNA aptamers in cells using autocatalytic transcripts. Nature biotechnology 37, 667–675 (2019).

39. Grubbs, R.D. Intracellular magnesium and magnesium buffering. Biometals : an international journal on the role of metal ions in biology, biochemistry, and medicine 15, 251–259 (2002).

40. Dai, L.J. & Quamme, G.A. Intracellular Mg2+ and magnesium depletion in isolated renal thick ascending limb cells. The Journal of clinical investigation 88, 1255–1264 (1991).

41. Novotny, I. et al. SART3-Dependent Accumulation of Incomplete Spliceosomal snRNPs in Cajal Bodies. Cell reports 10, 429–440 (2015).

42. Tourriere, H. et al. The RasGAP-associated endoribonuclease G3BP assembles stress granules. The Journal of cell biology 160, 823–831 (2003).

43. Sunbul, M. et al. Super-resolution RNA imaging using a rhodamine-binding aptamer with fast exchange kinetics. Nature biotechnology 39, 686–690 (2021).

44. Englert, D. et al. Fast-exchanging spirocyclic rhodamine probes for aptamer-based super-resolution RNA imaging. Nature communications 14, 3879 (2023).

45. Li, X. et al. Imaging Intracellular S-Adenosyl Methionine Dynamics in Live Mammalian Cells with a Genetically Encoded Red Fluorescent RNA-Based Sensor. Journal of the American Chemical Society 142, 14117–14124 (2020).

46. Wojciechowska, M. & Krzyzosiak, W.J. Cellular toxicity of expanded RNA repeats: focus on RNA foci. Human molecular genetics 20, 3811–3821 (2011).

47. Brook, J.D. et al. Molecular basis of myotonic dystrophy: expansion of a trinucleotide (CTG) repeat at the 3’ end of a transcript encoding a protein kinase family member. Cell 69, 385 (1992).

48. Ho, T.H. et al. Muscleblind proteins regulate alternative splicing. The EMBO journal 23, 3103–3112 (2004).

49. Batra, R. et al. Elimination of Toxic Microsatellite Repeat Expansion RNA by RNA-Targeting Cas9. Cell 170, 899–912 e810 (2017).

50. Ouyang, J.P.T., Shukla, S., Bensalah, M. & Parker, R. DM1 repeat-expanded RNAs confer RNA toxicity as individual nuclear-retained RNAs. Cell reports 44, 115582 (2025).

51. Hwang, D.W., Maekiniemi, A., Singer, R.H. & Sato, H. Real-time single-molecule imaging of transcriptional regulatory networks in living cells. Nature reviews. Genetics 25, 272–285 (2024).

52. Daigle, N. & Ellenberg, J. LambdaN-GFP: an RNA reporter system for live-cell imaging. Nature methods 4, 633–636 (2007).

53. Wu, J. et al. Live imaging of mRNA using RNA-stabilized fluorogenic proteins. Nature methods 16, 862–865 (2019).

54. Bertrand, E. et al. Localization of ASH1 mRNA particles in living yeast. Molecular cell 2, 437–445 (1998).

55. Li, X., Kim, H., Litke, J.L., Wu, J. & Jaffrey, S.R. Fluorophore-Promoted RNA Folding and Photostability Enables Imaging of Single Broccoli-Tagged mRNAs in Live Mammalian Cells. Angewandte Chemie 59, 4511–4518 (2020).

56. Buhler, B. et al. Avidity-based bright and photostable light-up aptamers for single-molecule mRNA imaging. Nature chemical biology 19, 478–487 (2023).

57. Voigt, F. et al. Single-Molecule Quantification of Translation-Dependent Association of mRNAs with the Endoplasmic Reticulum. Cell reports 21, 3740–3753 (2017).

58. M E Azzam & Algranati, I.D. Mechanism of Puromycin Action: Fate of Ribosomes after Release of Nascent Protein Chains from Polysomes. Proceedings of the National Academy of Sciences of the United States of America 70, 3866–3869 (1973).

59. Fang, M., Jiang, Y., Chen, X. & Yang, Y. Capabilities and challenges for the use of fluorescent RNAs in RNA dynamics research. Trends in cell biology 36, 86–99 (2026).

60. Lu, X., Kong, K.Y.S. & Unrau, P.J. Harmonizing the growing fluorogenic RNA aptamer toolbox for RNA detection and imaging. Chemical Society reviews 52, 4071–4098 (2023).

61. Zhou, W.J. et al. Fluorogenic Interacting Protein Stabilization for Orthogonal RNA Imaging. Angewandte Chemie, e202502350 (2025).

62. Kuffner, C.J., Marzilli, A.M. & Ngo, J.T. RNA-stabilized coat proteins for sensitive and simultaneous imaging of distinct single mRNAs in live cells. Nature methods 23, 153–164 (2026).

63. Pham, T.G. et al. Orthogonal RNA-regulated destabilization domains for three-color RNA imaging with minimal RNA perturbation. Nature methods 23, 165–174 (2026).

64. Xia, C., Colognori, D., Jiang, X.S., Xu, K. & Doudna, J.A. Single-molecule live-cell RNA imaging with CRISPR-Csm. Nature biotechnology 43, 2023–2030 (2025).

65. Xia, C., Colognori, D., Jiang, X.S., Xu, K. & Doudna, J.A. Single-molecule live-cell RNA imaging with CRISPR-Csm. Nature biotechnology (2025).

66. Tutucci, E. et al. An improved MS2 system for accurate reporting of the mRNA life cycle. Nature methods 15, 81–89 (2018).

67. Li, W., Maekiniemi, A., Sato, H., Osman, C. & Singer, R.H. An improved imaging system that corrects MS2-induced RNA destabilization. Nature methods 19, 1558–1562 (2022).

68. Garcia, J.F. & Parker, R. MS2 coat proteins bound to yeast mRNAs block 5’ to 3’ degradation and trap mRNA decay products: implications for the localization of mRNAs by MS2-MCP system. Rna 21, 1393–1395 (2015).

69. Haimovich, G. et al. Intercellular mRNA trafficking via membrane nanotube-like extensions in mammalian cells. Proceedings of the National Academy of Sciences of the United States of America 114, E9873–E9882 (2017).

70. Cawte, A.D., Unrau, P.J. & Rueda, D.S. Live cell imaging of single RNA molecules with fluorogenic Mango II arrays. Nature communications 11, 1283 (2020).

