## Supplementary information for "A superfolding sfPepper enables superresolution and single-molecule RNA imaging in live cells"

**Supplementary Figures**

**Figure S1**: The effects of inserting or deleting nucleotides in J3/2 on Pepper's fluorescence.

**Figure S2**: The effects of introducing a stem at different sites of J3/2 on Pepper's fluorescence.

**Figure S3**: The performance of Pepper variants in live mammalian cells.

**Figure S4**: Mutagenesis in Pepper for reducing its $Mg^{2+}$ dependence.

**Figure S5**: Cellular performance of Pepper variants generated through combinatorial mutagenesis.

**Figure S6**: Comparison of sfPepper and Pepper in live cells incubated with different HBC analogues.

**Figure S7**: In vitro characteristics of sfPepper.

**Figure S8**: Comparison of the cellular brightness of sfPepper and Pepper alone or embedded in a tRNA scaffold.

**Figure S9**: Imaging of sfPepper-tagged small noncoding RNAs.

**Figure S10**: Tandem repeats of sfPepper aptamer.

**Figure S11**: Imaging of stress granules using sfPepper.

**Figure S12**: Comparison of sfPepper to other high-performance FRs.

**Figure S13**: Imaging of CUG repeat-containing RNA using different FRs.

**Figure S14**: Cotranslational localization of *Srprb* mRNA to the ER.

**Figure S15:** Single-molecule RNA imaging using 4sfPepper.

**Figure S16**: Comparison of the brightness of 4sfPepper- and 4×4sfPepper-tagged single-molecule mRNAs.

**Figure S17**: Single-molecule RNA imaging using four repeats of Pepper or RhoBAST aptamer.

**Figure S18**: Real-time tracking of single-molecule mRNAs in live cells.

**Figure S19**: Analysis of the diffusion coefficients for different ER-localized mRNA transcripts.

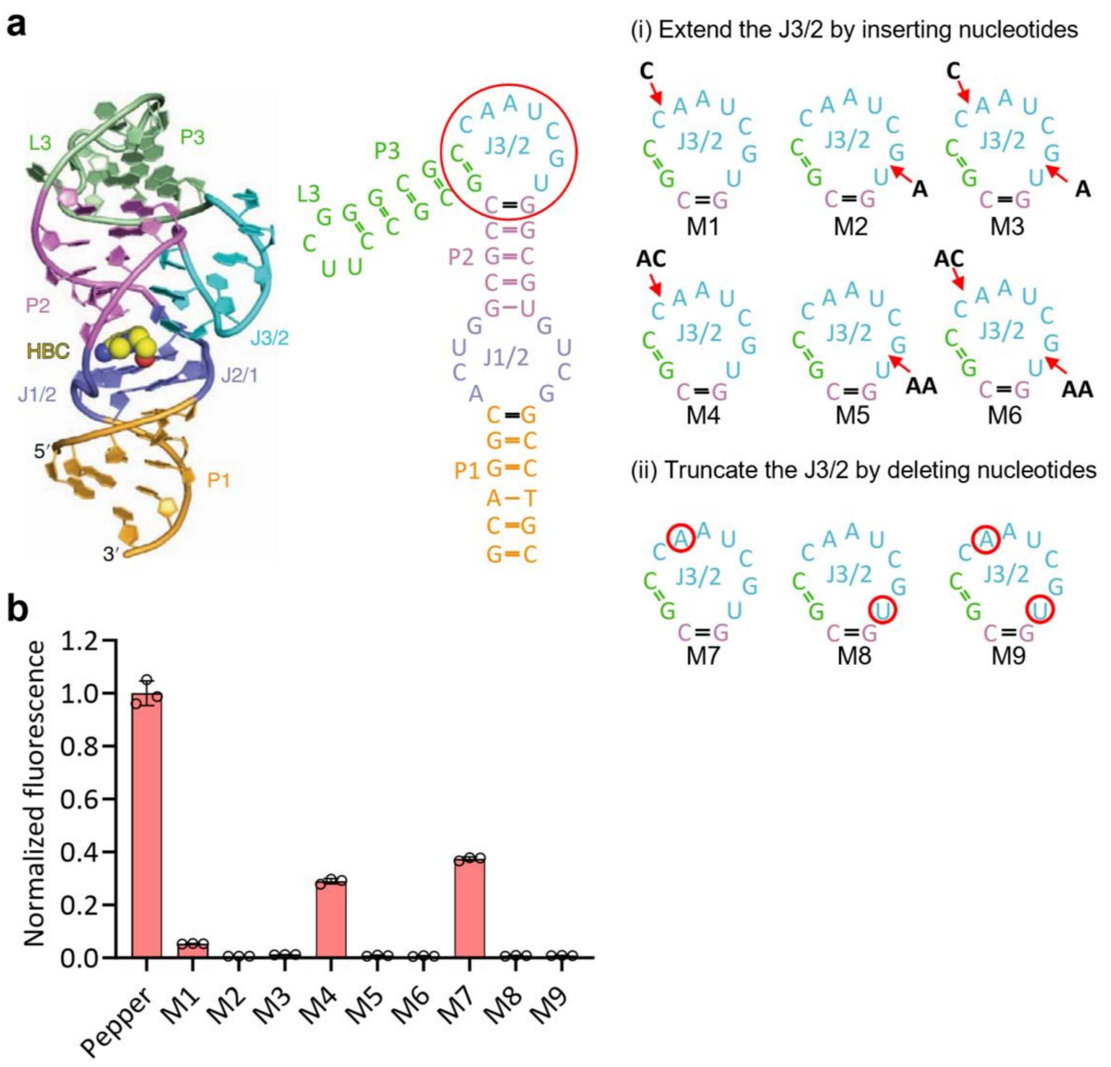


**Supplementary Figure 1 The effects of inserting or deleting nucleotides in J3/2 on Pepper's fluorescence. (a)** Tertiary and secondary structures of Pepper (PDB: 7EOK). The nucleotides for insertion or deletion are indicated with red arrows and circles, respectively. The left panel of tertiary structure is adapted from ref. 11. **(b)** The capability of different Pepper variants to activate the fluorescence of HBC620. Data were normalized to the fluorescence of original Pepper, respectively. The data represent the means ± SDs from three biologically independent replicates.

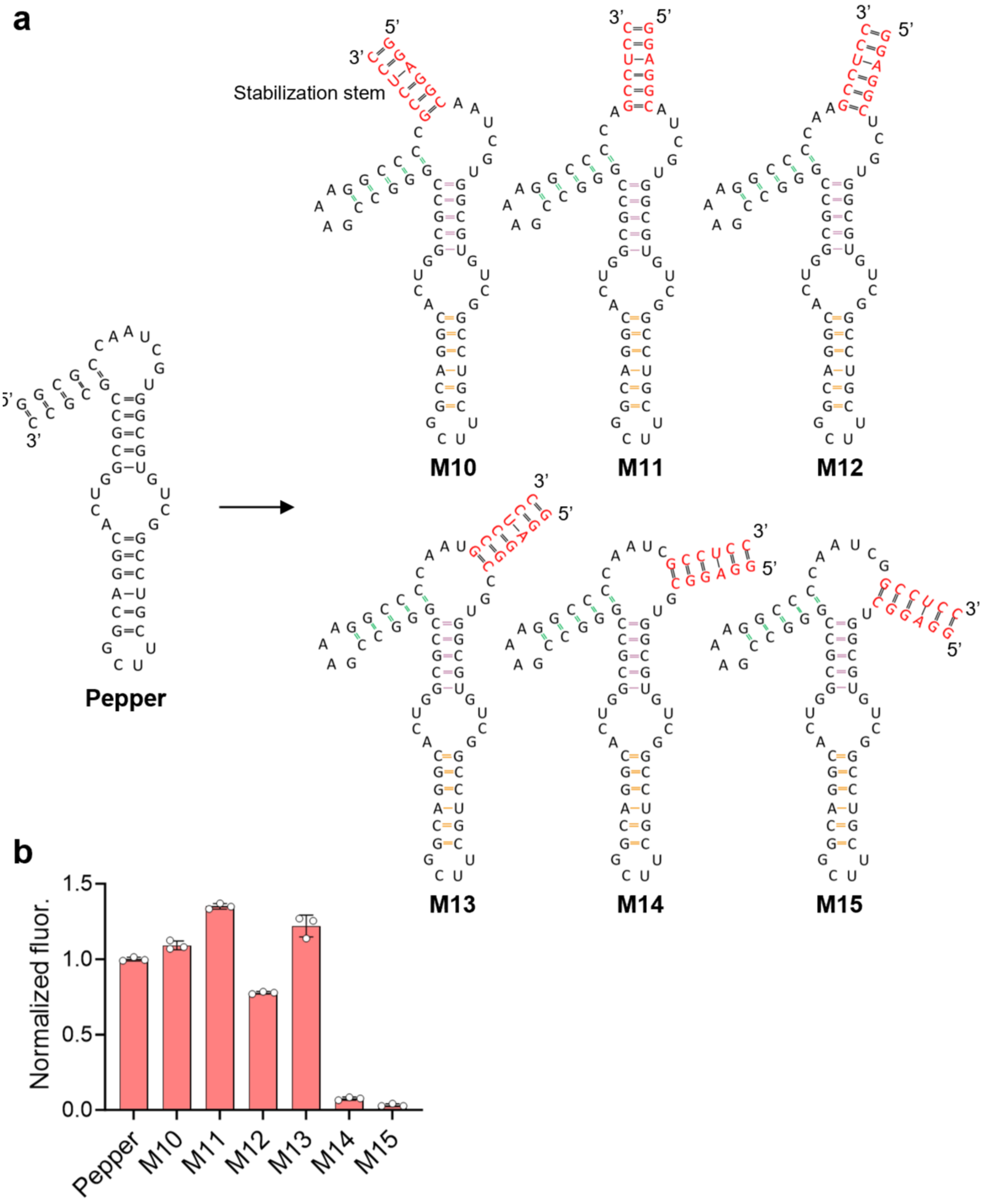


**Supplementary Figure 2 The effects of introducing a stem at different sites of J3/2 on Pepper's fluorescence. (a)** Schematic diagram of inserting a stem at different sites of J3/2 in Pepper. **(b)** The capability of different Pepper variants to activate the fluorescence of HBC620. Data were normalized to the fluorescence of original Pepper, respectively. The data represent the means ± SDs from three biologically independent replicates.

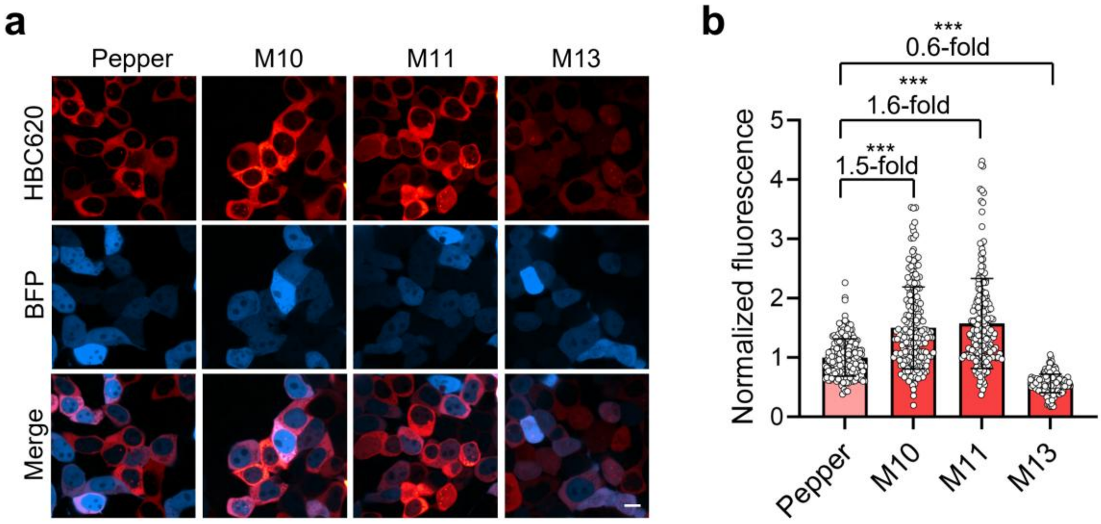


**Supplementary Figure 3 The performance of Pepper variants in live mammalian cells. (a)** Fluorescence imaging of live HEK293T cells expressing different Pepper variants upon incubation with 0.5 μM HBC620 dye. The cells were cotransfected with a plasmid expressing blue fluorescent protein TagBFP to distinguish transfected cells from nontransfected ones. Scale bar, 10 μm. **(b)** The fluorescence intensities of the cells in **a**. Statistical comparisons were performed by two-tailed *t* tests. ***$P < 0.001$. The data represent the means ± SDs ($N = 200$ cells).

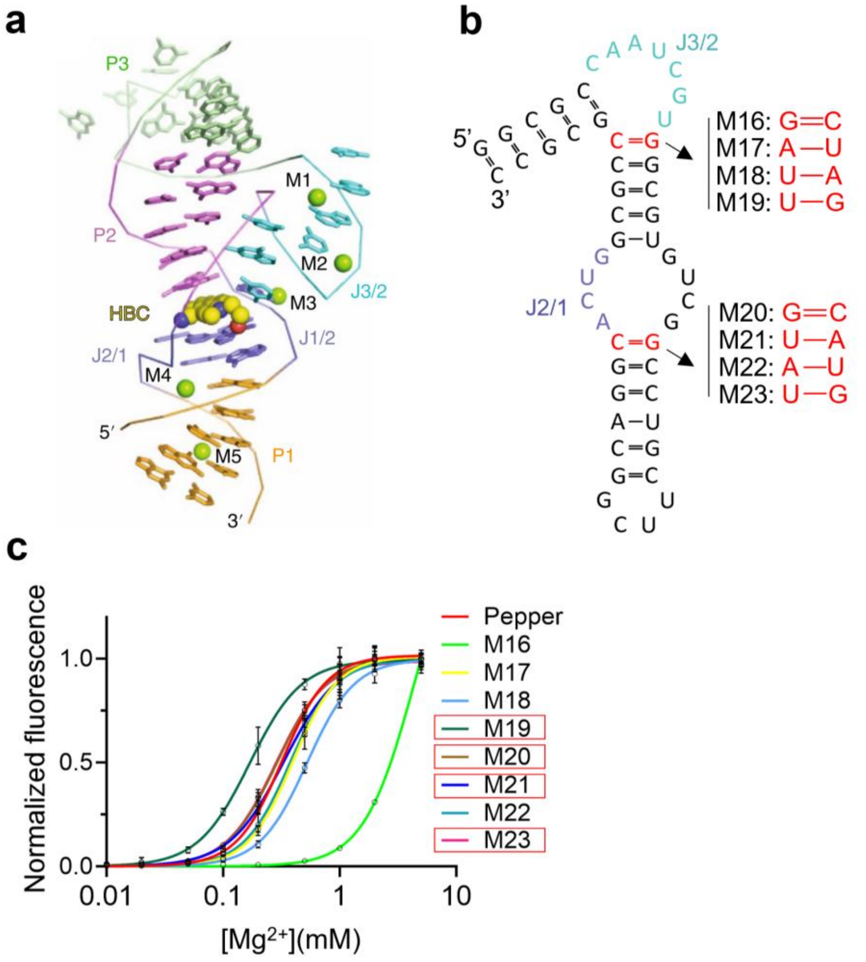


**Supplementary Figure 4 Mutagenesis in Pepper for reducing its $Mg^{2+}$ dependence. (a)** Schematic diagram of five divalent metal cations (M1–M5) identified in the tertiary structure of Pepper aptamer in complex with HBC, four of which (M1–M4) are located in the junction regions J3/2 and J2/1, and one of which (M5) is bound to the stem P1. The figure is adapted from Ref 11. **(b)** Targeted mutations at conformational junctions, specifically, the nucleotide pairs bridging J3/2 and P2, and J2/1 and P1 stem, with the aim of reducing $Mg^{2+}$ dependence without compromising Pepper's structural integrity. **(c)** $Mg^{2+}$ dependence of different Pepper variants in **b**. 0.2 µM RNA aptamer was mixed with 2 µM HBC620, and the fluorescence signal of the complex was measured with different concentrations of $MgCl_2$. Variants with reduced $Mg^{2+}$ dependence compared to Pepper were indicated by red boxes. Data represent the means ± SDs from three biologically independent replicates.

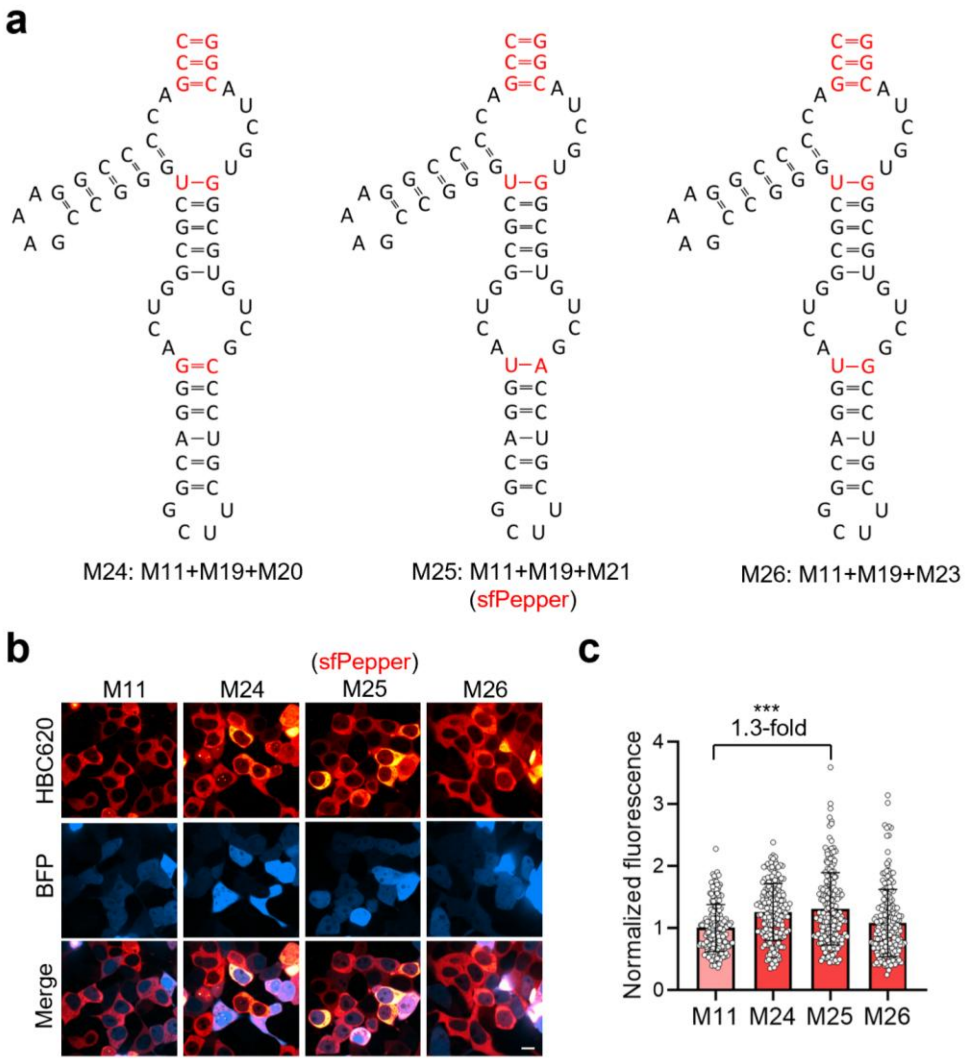


**Supplementary Figure 5 Cellular performance of Pepper variants generated through combinatorial mutagenesis. (a)** Schematic diagram of Pepper variants generated through combinatorial mutagenesis. **(b)** The performance of different Pepper variants in live mammalian cells. HEK293T cells expressing different Pepper variants were incubated with 0.5 μM HBC620 and imaged. The cells were cotransfected with a plasmid expressing blue fluorescent protein TagBFP to distinguish transfected cells from nontransfected ones. Scale bar, 10 μm. **(c)** The fluorescence intensities of the cells in **b**. Statistical comparison was performed by a two-tailed *t* test. ***$P < 0.001$. The data represent the means ± SDs ($N = 200$ cells).

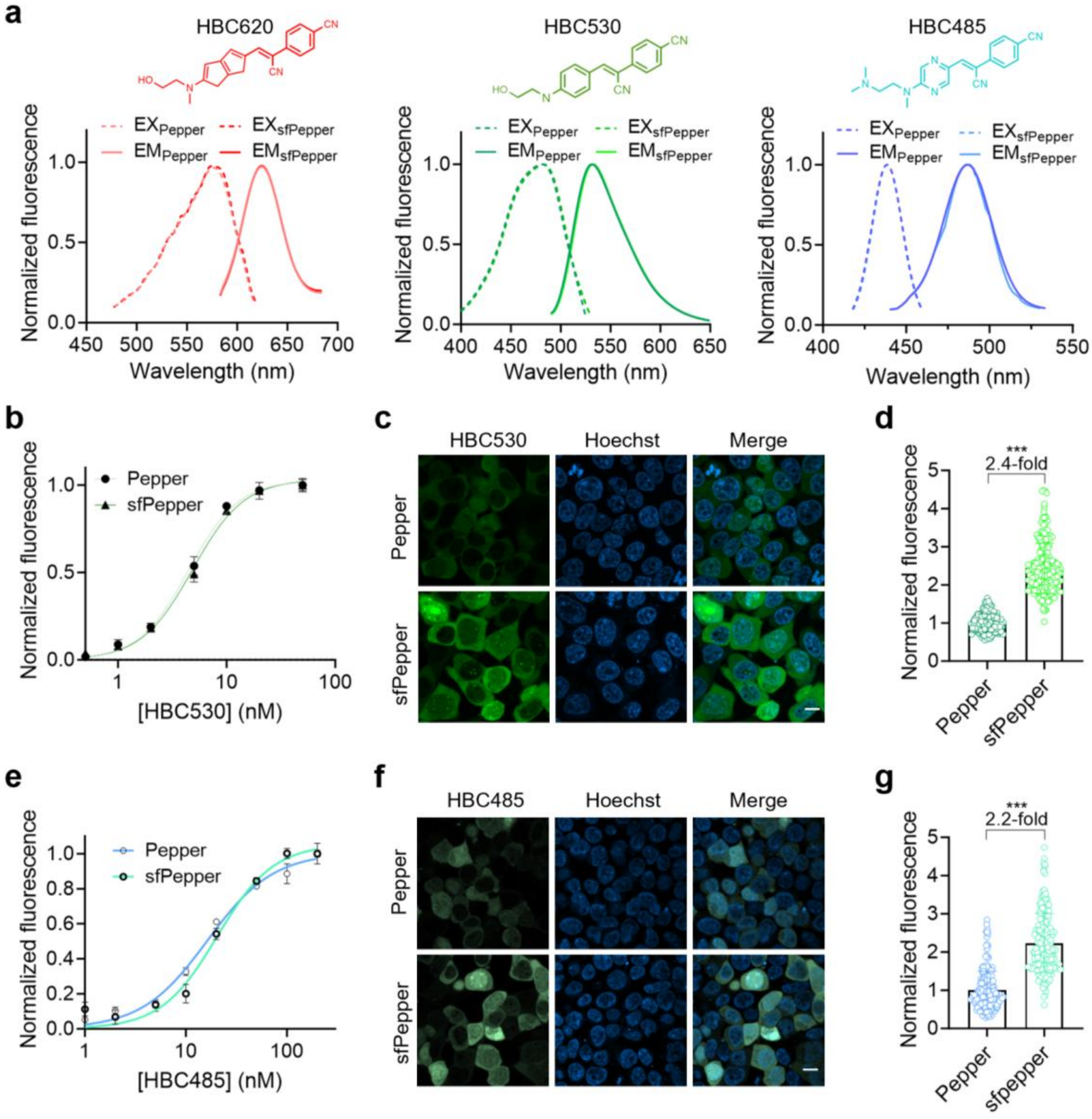


**Supplementary Figure 6 Comparison of sfPepper and Pepper in live cells incubated with different HBC analogues. (a)** The excitation and emission spectra of Pepper and sfPepper upon incubation with different analogues. **(b)** Binding affinity of HBC530 to Pepper or sfPepper. The fluorescence of RNA-dye complexes in the presence of increasing concentrations of HBC530 was measured. Data represent the means ± SDs from three biologically independent replicates. **(c)** Confocal images of live HEK293T cells expressing circular Pepper or sfPepper incubated with 0.5 µM HBC530. Scale bar, 10 µm. **(d)** The fluorescence intensities of the cells in **c**. Statistical comparison was performed by a two-tailed *t* test. ***$P < 0.001$. The data represent the means ± SDs ($N = 200$ cells). **(e)** Binding affinity of HBC485 to Pepper or sfPepper. The fluorescence of RNA-dye complexes in the presence of increasing concentrations of HBC485 was measured. **(f)** Confocal images of live HEK293T cells expressing circular Pepper or sfPepper incubated with 0.5 µM HBC485. Scale bar, 10 µm. **(g)** The fluorescence intensities of the cells in **f**. Statistical comparison was performed by a two-tailed *t* test. ***$P < 0.001$. The data represent the means ± SDs ($N$ = 200 cells).

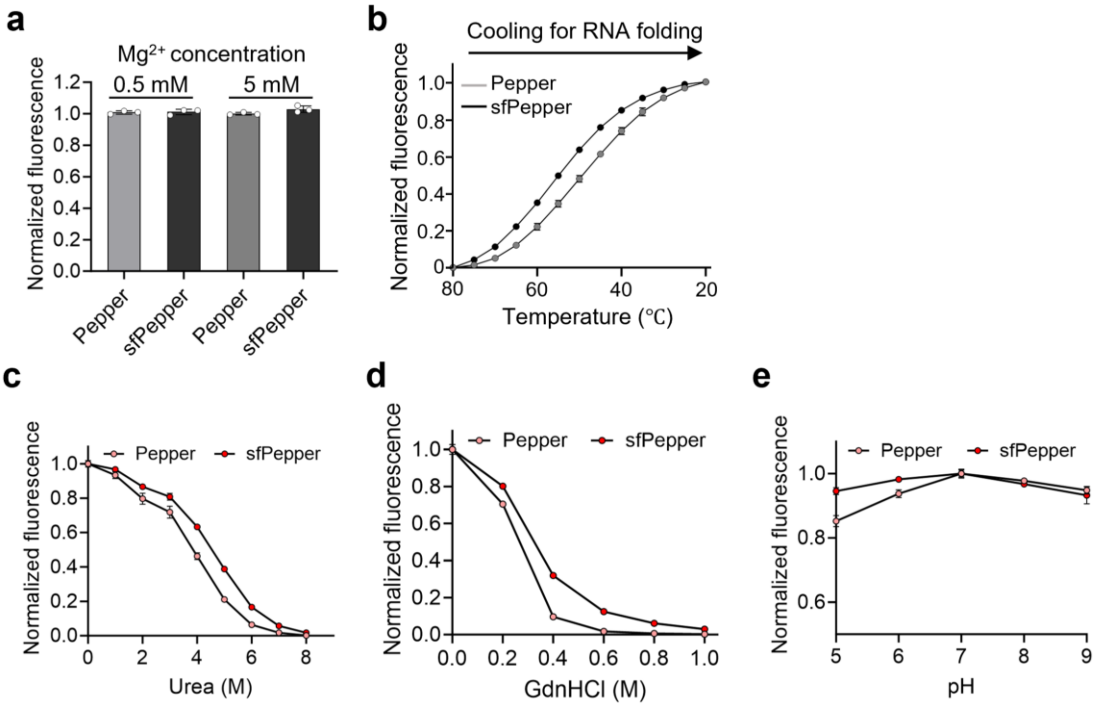


**Supplementary Figure 7 In vitro characteristics of sfPepper. (a)** The fluorescence intensities of Pepper and sfPepper in the presence of excess RNA aptamer and 0.5 or 5 mM $Mg^{2+}$. **(b)** Folding kinetics of sfPepper and Pepper. The RNA was denatured at 80°C for 10 min and then cooled to 20 °C. The fluorescence of the complexes at the indicated temperatures was measured. **(c)** Fluorescence intensity of sfPepper and Pepper under varying urea concentrations. **(d)** Fluorescence intensity of sfPepper and Pepper under varying GdnHCl concentrations. **(e)** Fluorescence intensity of sfPepper and Pepper across a pH range of 5 to 9. The data in **a-e** represent the means ± SDs from three biologically independent replicates.

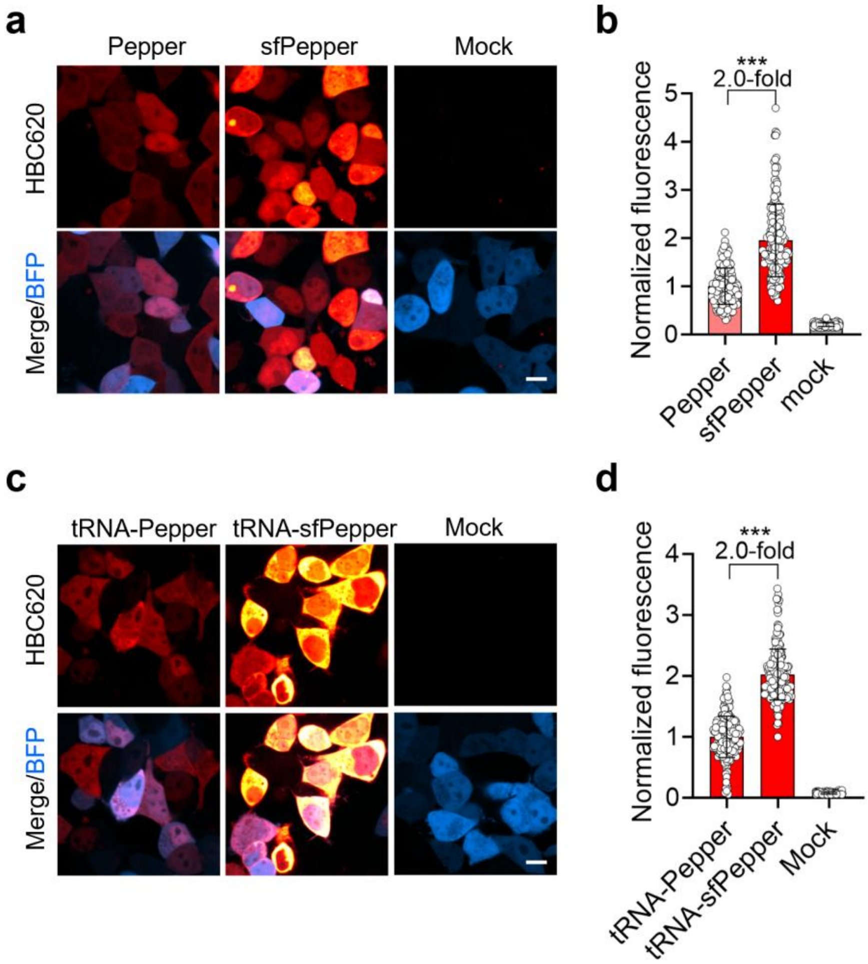


**Supplementary Figure 8 Comparison of the cellular brightness of sfPepper and Pepper alone or embedded in a tRNA scaffold. (a)** Confocal images of HEK293T cells expressing linearized sfPepper or Pepper incubated with 0.5 µM HBC620. The cells were cotransfected with a plasmid expressing blue fluorescent protein TagBFP to distinguish transfected cells from nontransfected ones. The cells expressing TagBFP alone were used as the control. Scale bar, 10 µm. **(b)** The fluorescence intensities of the cells in **a**. Statistical comparison was performed by a two-tailed *t* test. ****P* < 0.001. Data represent the means ± SDs (*N*= 200 cells). **(c)** Confocal images of HEK293T cells expressing sfPepper or Pepper embedded in a tRNA scaffold incubated with 0.5 µM HBC620. The cells were cotransfected with a plasmid expressing blue fluorescent protein TagBFP to distinguish transfected cells from nontransfected ones. The cells expressing TagBFP alone were used as the control. Scale bar, 10 µm. **(d)** The fluorescence intensities of the cells in **c**. Statistical comparison was performed by a two-tailed *t* test. ****P* < 0.001. Data represent the means ± SDs (*N* = 200 cells).

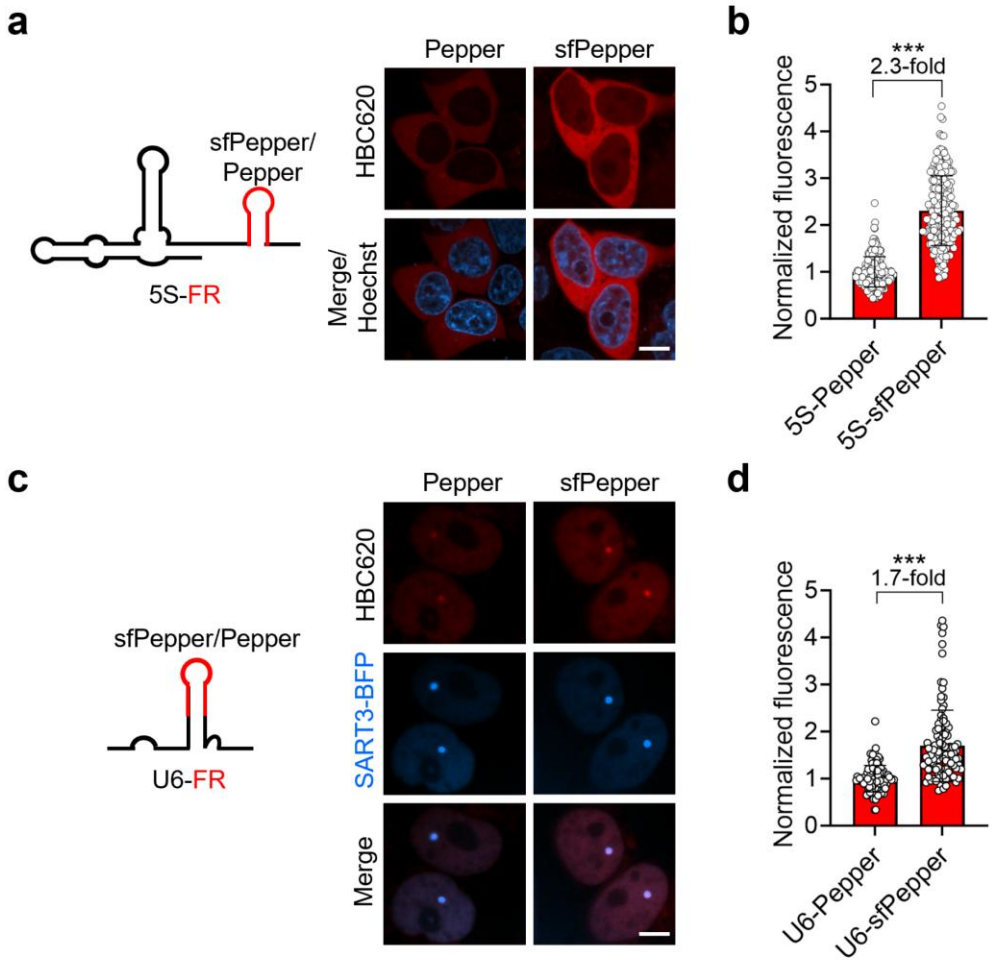


**Supplementary Figure 9 Imaging of sfPepper-tagged small noncoding RNAs. (a)** Confocal images of HEK293T cells expressing sfPepper- or Pepper-tagged 5S ribosomal RNA. Scale bar, 10 µm. **(b)** The fluorescence intensities of the cells in **a**. Statistical comparison was performed by a two-tailed *t* test. $^{***}P < 0.001$. The data represent the means ± SDs ($N = 200$ cells). **(c)** Confocal images of HEK293T cells expressing sfPepper- or Pepper-tagged U6 nuclear RNA. The cells were co-transfected with a plasmid expressing BFP-tagged SART3 serving as the marker for Cajar bodies. Scale bar, 5 µm. **(d)** The fluorescence intensities of the cells in **c**. Statistical comparison was performed by a two-tailed *t* test. $^{***}P < 0.001$. The data represent the means ± SDs ($N = 110$ cells).

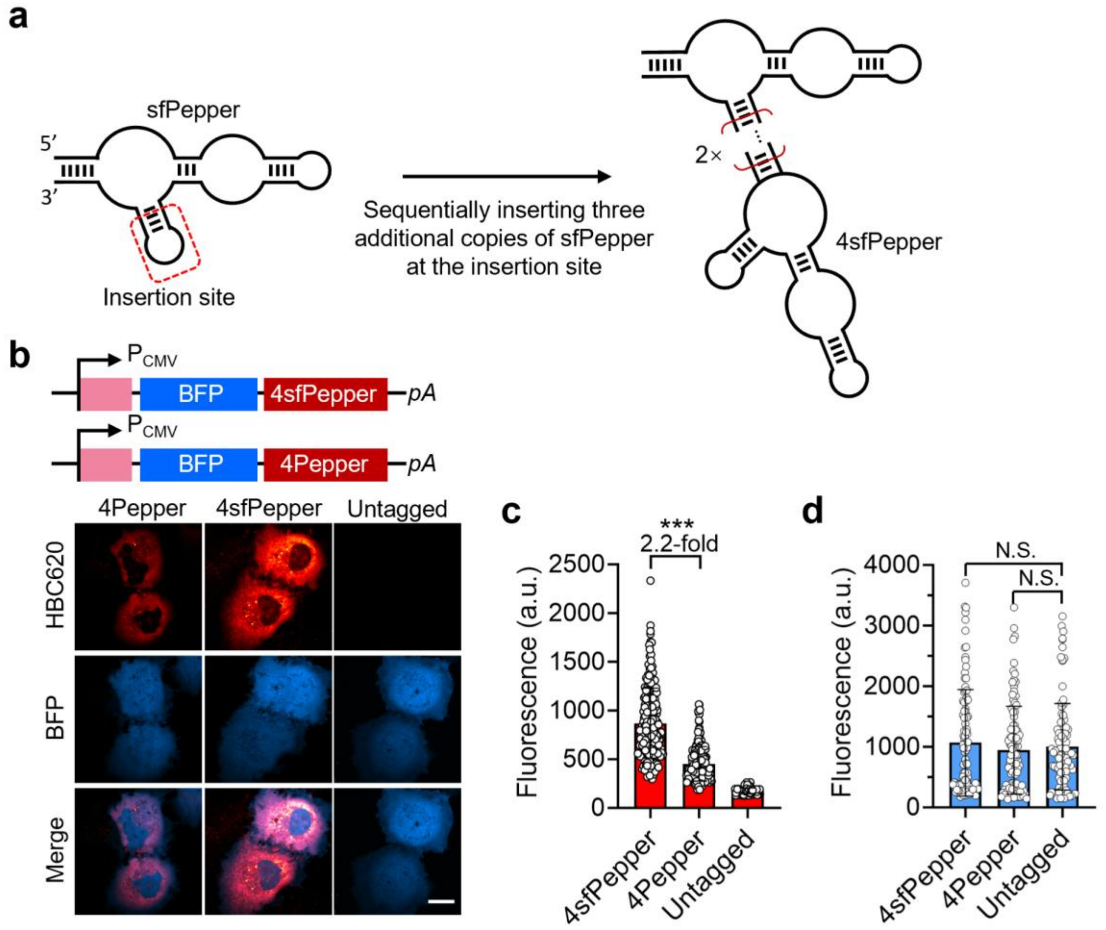


**Supplementary Figure 10 Tandem repeats of sfPepper aptamer. (a)** Schematic representation of four tandem repeat of sfPepper aptamer. One molecule sfPepper is inserted into the stem-loop of another molecule sfPepper. **(b)** Confocal images of COS-7 cells expressing 4sfPepper- or 4Pepper-tagged BFP mRNA upon incubation with 0.2 μM HBC620. The cells expressing untagged BFP mRNA were used as the control. Scale bar, 10 μm. **(c, d)** Quantitative analysis of red **(c)** and blue **(d)** fluorescence in individual cells in **b**. Statistical comparisons were performed by two-tailed *t* tests. ***$P < 0.001$. N.S., no significant difference. The data represent the means ± SDs ($N = 100$ cells).

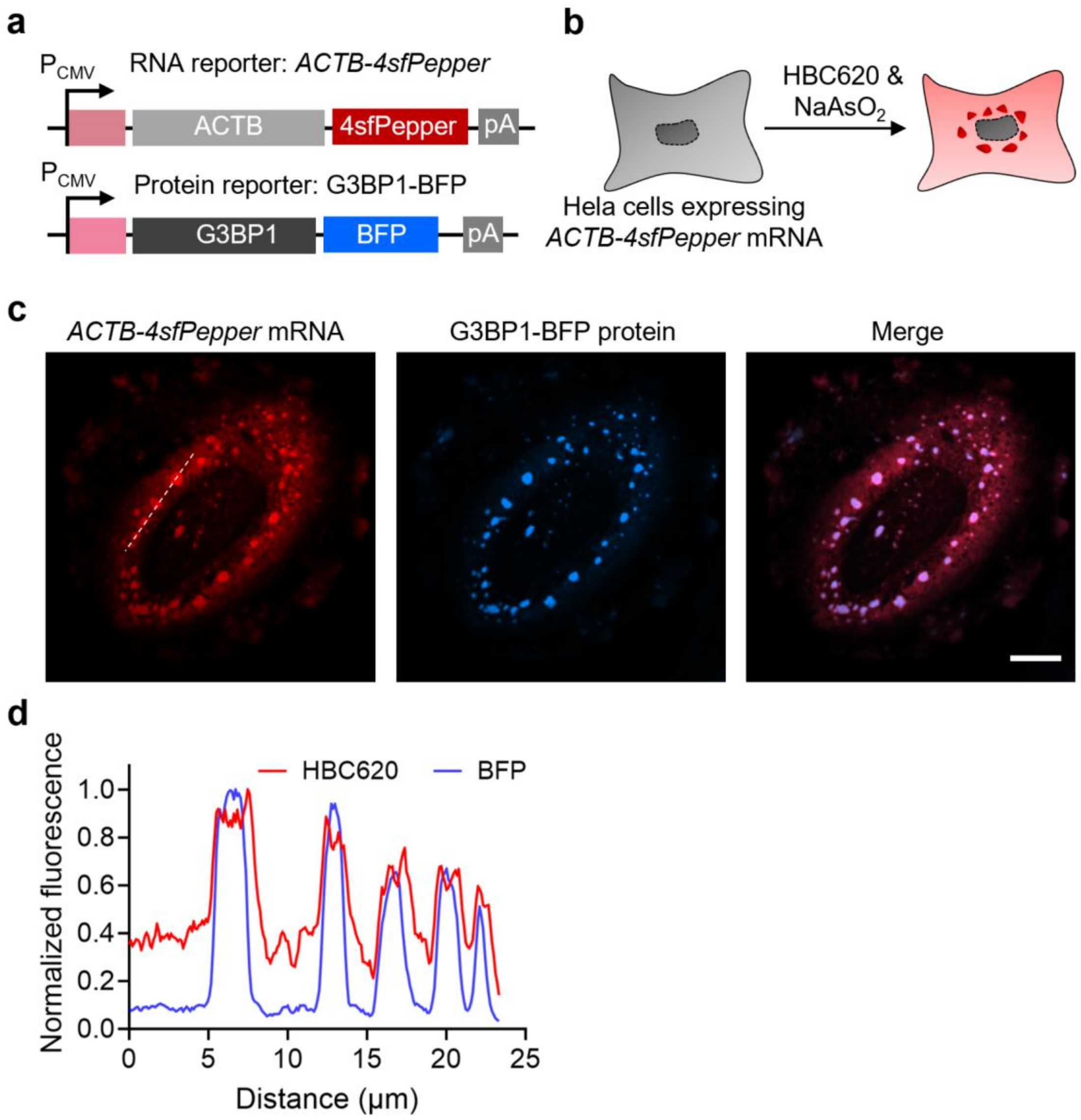


**Supplementary Figure 11 Imaging of stress granules using sfPepper. (a)** Schematic representation of constructs expressing 4sfPepper-tagged *ACTB* mRNA and G3BP1-BFP fusion protein. G3BP1 protein is a granule-associated RNA-binding protein that can serve as a marker to localize stress granules (SGs). **(b)** Schematic illustration of sfPepper-based imaging of SGs. After transient transfection of plasmids expressing ACTB-4sfPepper and G3BP1-BFP, SGs were induced by sodium arsenite ($NaAsO_2$) treatment and imaged. **(c)** Imaging of ACTB-4sfPepper and G3BP1-BFP in cells treated with 0.2 μM HBC620 and 200 μM $NaAsO_2$. Scale bar, 10 μm. **(d)** Normalized intensity profiles along the white line in **c**.

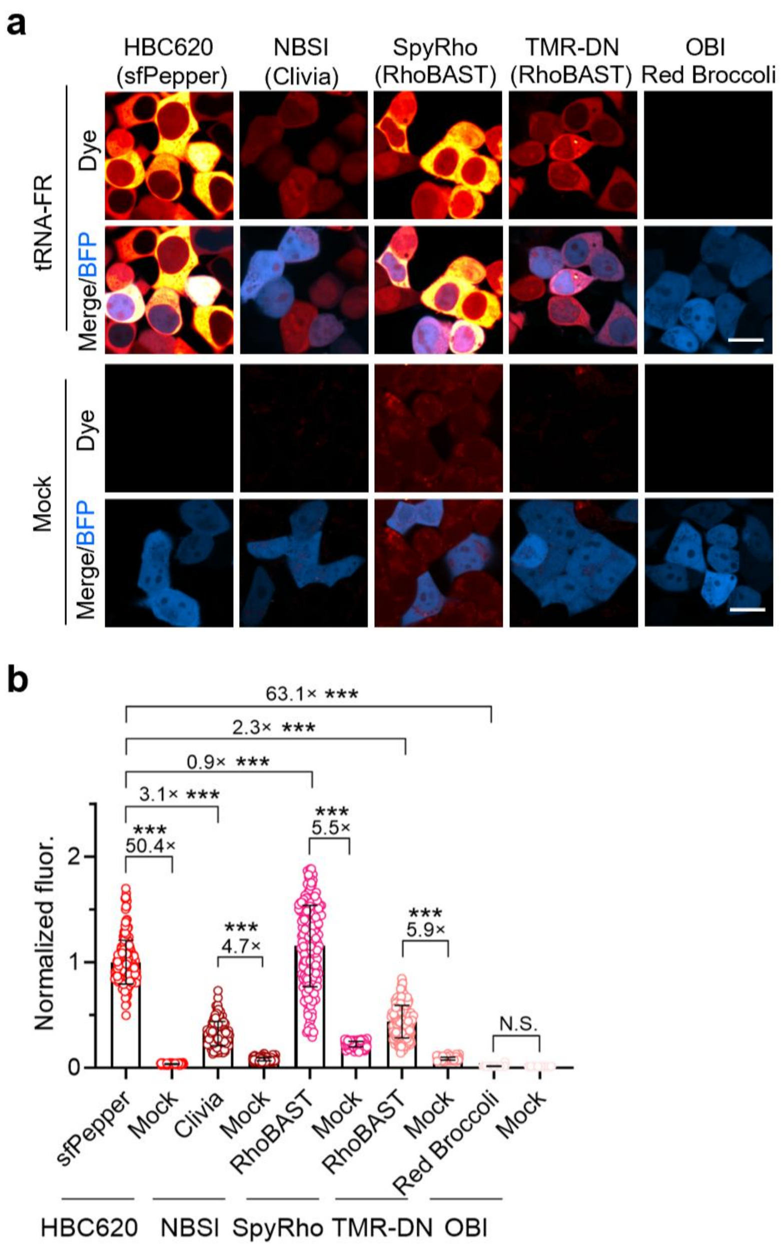


**Supplementary Figure 12 Comparison of sfPepper to other high-performance FRs. (a)** Confocal images of HEK293T cells expressing sfPepper or Pepper embedded in a tRNA scaffold. The fluorescence was imaged by incubation with HBC620 (0.5 µM), NBSI (0.5 µM), SpyRho (0.5 µM), TMR-DN (0.5 µM), and OBI (0.5 µM), respectively. The cells were cotransfected with a plasmid expressing blue fluorescent protein TagBFP to distinguish transfected cells from nontransfected ones. The control cells were transfected with a plasmid expressing TagBFP alone and incubated with the respective dye. Scale bars, 10 µm. **(b)** The fluorescence intensities of the cells in **a**. Statistical comparisons were performed by two-tailed *t* tests. ***$P < 0.001$. The data represent the means ± SDs ($N$ = 200 cells).

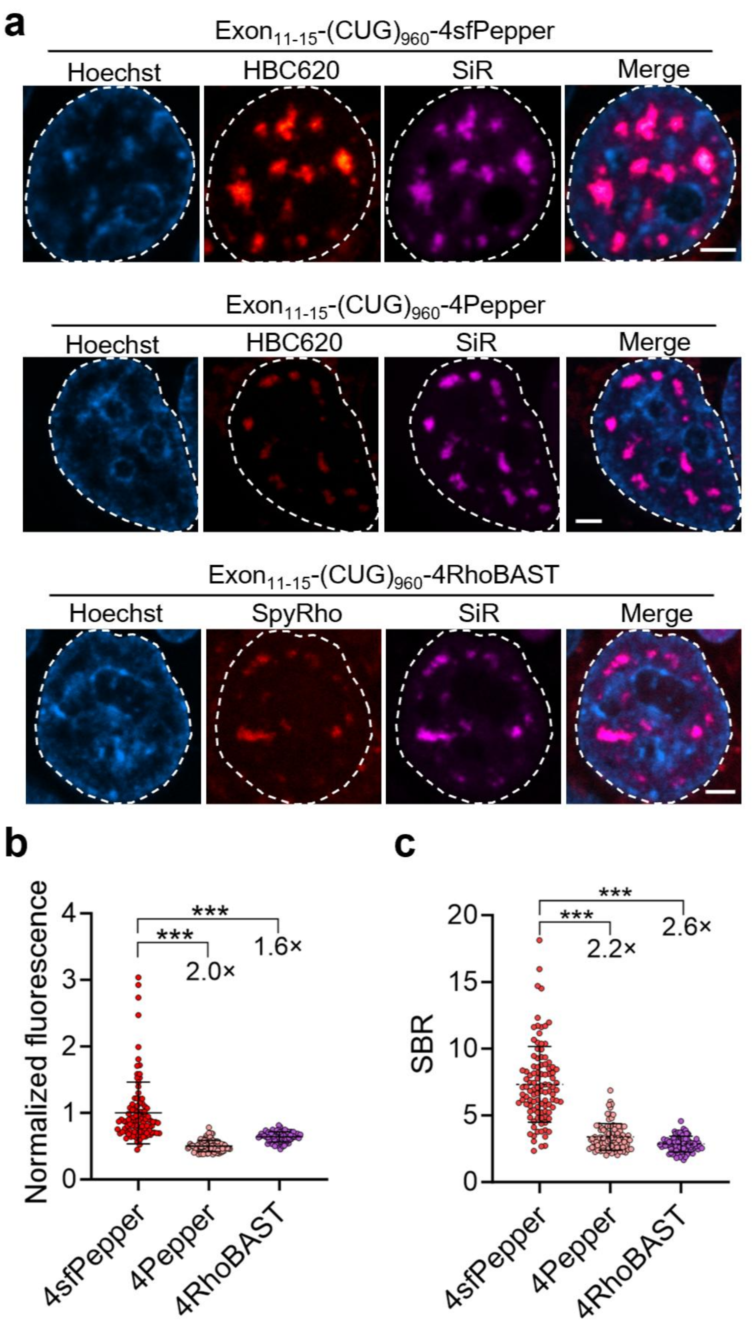


**Supplementary Figure 13 Imaging of CUG repeat-containing RNA using different FRs. (a)** Confocal images of HEK293T cells expressing 4sfPepper-, 4Pepper- or 4RhoBAST-tagged CUG repeat RNA upon incubation with 0.5 µM HBC620 (4sfPepper and 4Pepper) or SpyRho (4RhoBAST), respectively. Scale bars, 5 µm. **(b, c)** The fluorescence intensities **(b)** and signal-to-noise ratios **(c)** of the nuclear foci in **a**. The signal-to-noise ratios of the nuclear foci were calculated by dividing the fluorescence intensity of the foci by that of the nucleoplasm. Statistical comparisons were performed by two-tailed *t* tests. ***$P < 0.001$. The data represent the means ± SDs ($N$ = 104, 86 and 80 cells for 4sfPepper, 4Pepper and 4RhoBAST, respectively).

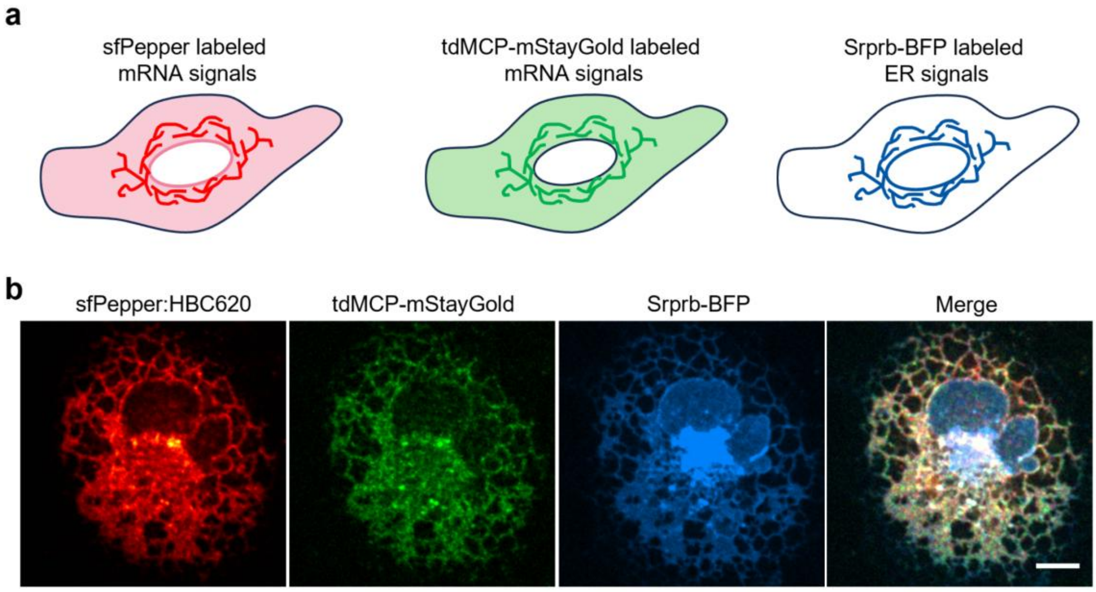


**Supplementary Figure 14 Cotranslational localization of Srprb mRNA to the ER. (a)** Schematic representation of expected ER-localized of *Srprb* mRNA simultaneously tagged by sfPepper and the MS2-MCP system, as well as the ER-localized Srprb-BFP fusion protein. **(b)** Confocal images of live COS-7 cells co-expressing *Srprb-BFP-4×4sfPepper-16×MS2* mRNA and tdMCP-mStayGold fusion protein upon incubation with 100 nM HBC620. Scale bar, 5 µm.

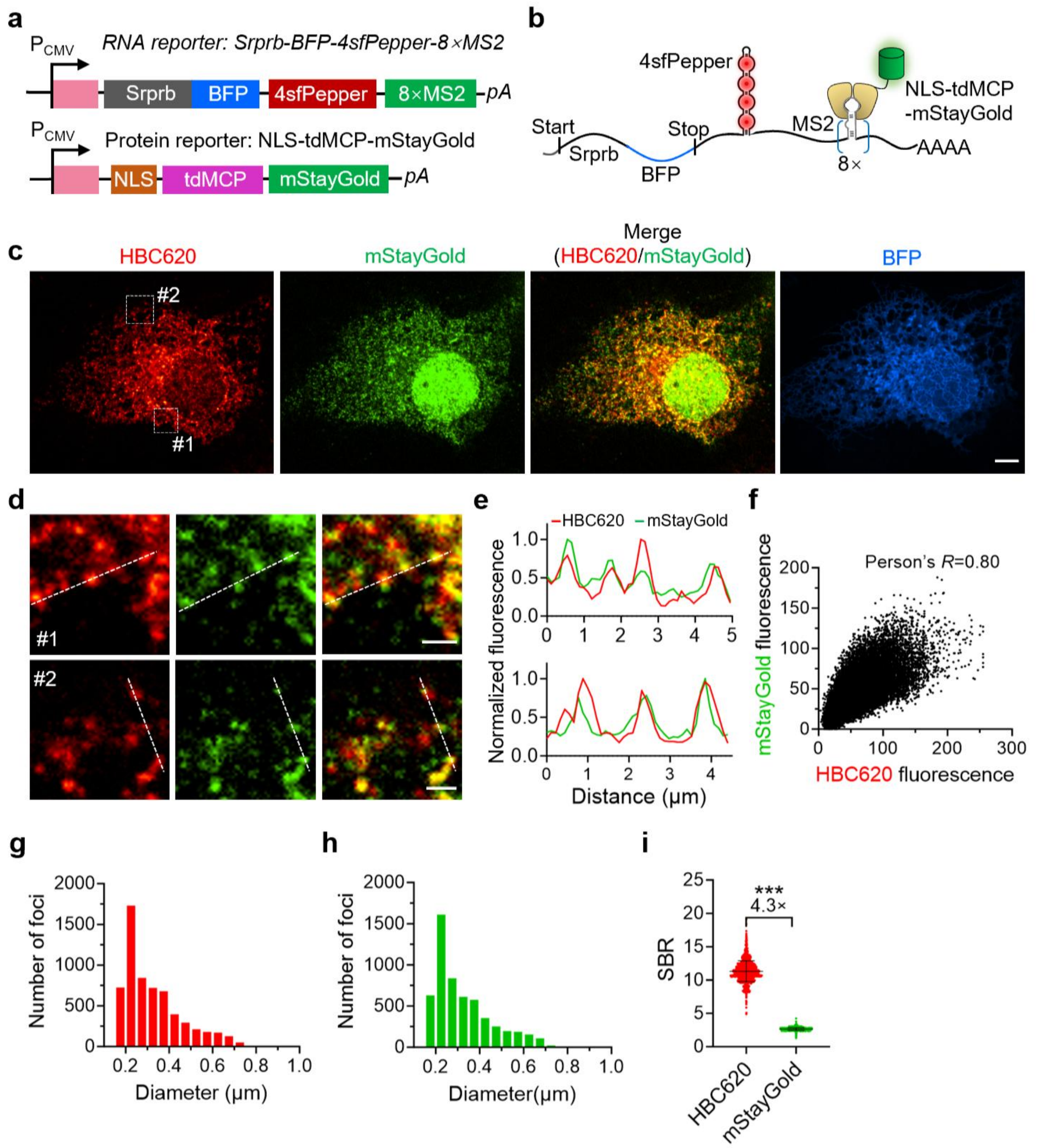


**Supplementary Figure 15 Single-molecule RNA imaging using 4sfPepper. (a)** Constructs used to express with 4Pepper- and 16×MS2-tagged *Srprb-BFP* mRNA, which can be bound by the fusion protein of tandem dimer of MCP (tdMCP) and mStayGold. **(b)** Illustration of *Srprb-BFP* mRNA tagged with 4Pepper and 16×MS2. **(c)** Confocal images of live COS-7 cells expressing low levels of *Srprb-BFP-4sfPepper-16×MS2* mRNA and tdMCP-mStayGold upon incubation with 100 nM HBC620. Scale bar, 10 µm. **(d)** Magnified views of the white box indicated regions in **(c)** to show the colocalization of red and green foci. Scale bars, 2 µm. **(e)** Line profiles of the fluorescence from HBC620 and mStayGold channels of the dashed lines in **d**. **(f)** Colocalization analysis of the cytosolic HBC620 and mStayGold signals in **c** (Pearson's $R$ = 0.80). **(g, h)** Analysis of the foci (*N* = 6,128 foci) in the HBC620 and (*N* = 5,545 foci) GFP channel as described in panels **d**–**f. (i)** The average signal-to-background ratio of foci in **c** from HBC620 and mStayGold channels. The data represent means ± SDs (HBC620: *N* = 1,905 foci; mStayGold: *N* = 3,113 foci). Statistical comparison was performed by a two-tailed *t* test. ***$P < 0.001$.

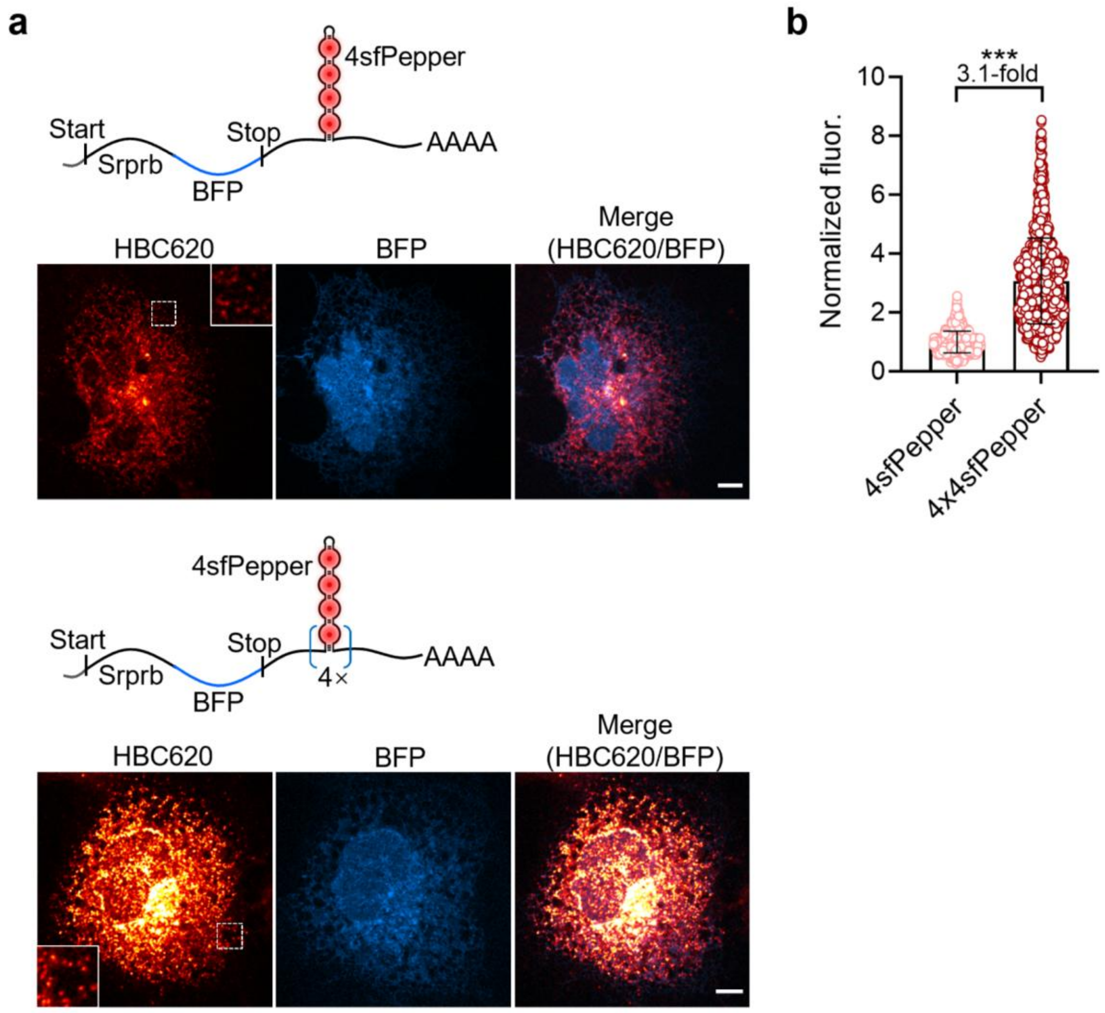


**Supplementary Figure 16 Comparison of the brightness of 4sfPepper- and 4×4sfPepper-tagged single-molecule mRNAs. (a)** Confocal images of COS-7 cells expressing *Srprb-BFP-4sfPeppe* or *Srprb-BFP-4×4sfPepper* mRNA incubated with 0.1 µM HBC620. Insets, magnified views of the white box indicated regions in the figures. Scale bars, 10 µm. **(b)** Quantitative analysis of the fluorescence intensity of individual single-molecule mRNA molecules in **a**. Statistical comparison was performed by a two-tailed *t* test. ****P* < 0.001. The data represent the means ± SDs (*N* = 1,590 particles).

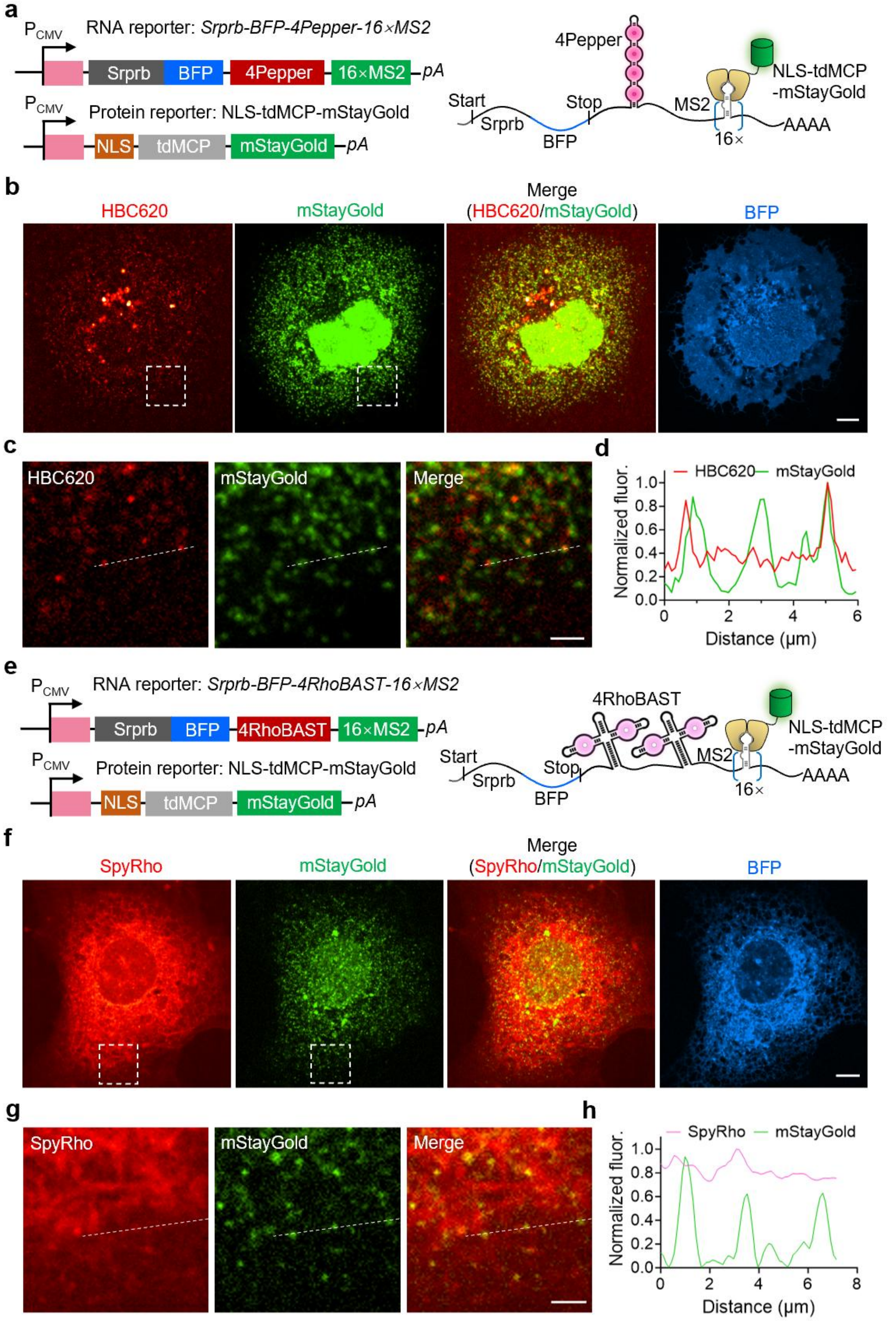


**Supplementary Figure 17 Single-molecule RNA imaging using four repeats of Pepper or RhoBAST aptamer. (a)** Constructs used to express with 4Pepper- and 16×MS2-tagged *Srprb-*

*BFP* mRNA, which can be bound by the fusion protein of tandem dimer of MCP (tdMCP) and mStayGold. **(b)** Confocal images of COS-7 cells expressing *Srprb-BFP-4Pepper-16×MS2* mRNA and tdMCP-mStayGold incubated with 0.1 µM HBC620. Scale bar, 10 µm. **(c)** Magnified views of the white box indicated regions in **b** to show the colocalization of red and green foci. Scale bar, 2 µm. **(d)** Line profiles of the fluorescence from HBC620 and mStayGold channels of the dashed lines in **c**. **(e)** Constructs used to express *Srprb-BFP-4RhoBAST-16×MS2* mRNA, which can be bound by tdMCP-mStayGold fusion protein. **(f)** Confocal images of COS-7 cells expressing *Srprb-BFP-4RhoBAST-16×MS2* mRNA and tdMCP-mStayGold protein incubated with 0.1 µM SpyRho. Scale bar, 10 µm. **(g)** Magnified views of the white box indicated regions in **f** to show the colocalization of red and green foci. Scale bar, 2 µm. **(h)** Line profiles of the fluorescence from HBC620 and mStayGold channels of the dashed lines in **g**.

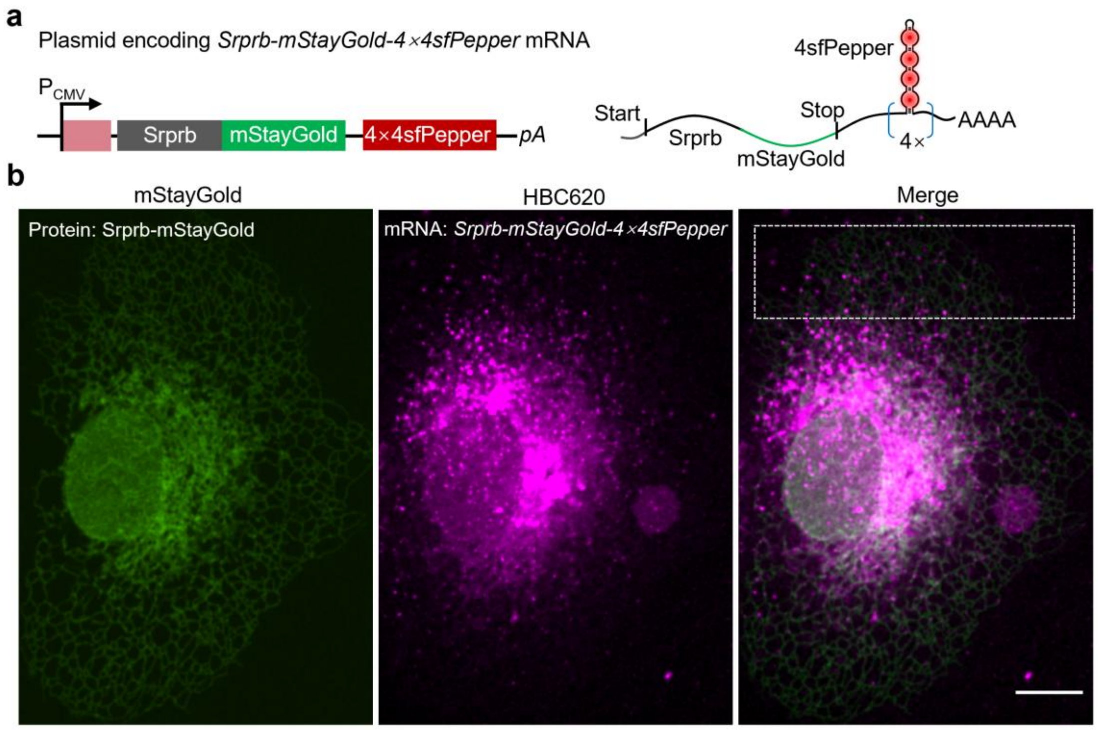


**Supplementary Figure 18 Real-time tracking of single-molecule mRNAs in live cells. (a)** Schematic representation of 4×4sfPepper-tagged *Srprb-mStayGold* mRNA. **(b)** Confocal images of a COS-7 cell expressing *Srprb-mStayGold-4×4sfPepper* mRNA incubated with 200 nM HBC620. Magnified view of the white box indicated region was shown in **Figure 4j**. Scale bar, 10 µm.

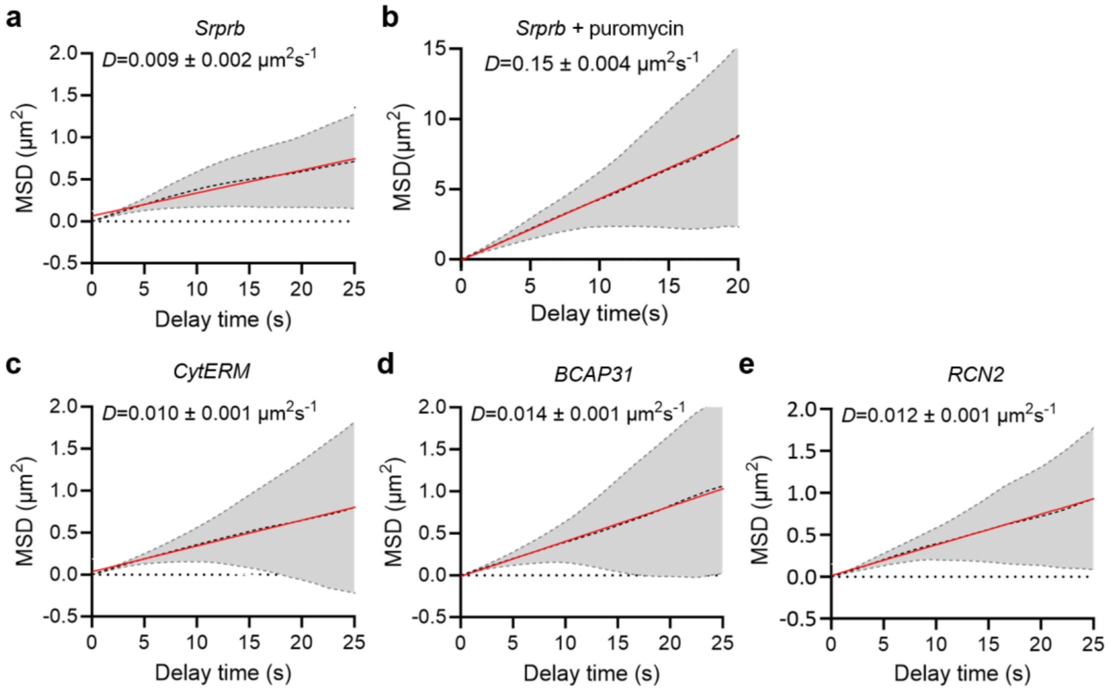


**Supplementary Figure 19 Analysis of the diffusion coefficients for different ER-localized mRNA transcripts.** COS-7 cells expressing *Srprb-mStayGold-4×4Pepper* **(a, b)**, *CytERM-BFP-4×4Pepper* **(c)**, *BCAP31-BFP-4×4Pepper* **(d),** or *RCN2-BFP-4×4Pepper* **(e)** mRNA were incubated with 0.2 μM HBC620 and imaged. For **b**, the cells were treated with 100 μg/mL puromycin prior to imaging. The diffusion coefficients for different mRNA transcripts were analyzed (*N* = 46, 46, 35, 50 and 33 tracks for **a** to **e**, respectively).
